# Wall teichoic acids regulate peptidoglycan synthesis by paving cell wall nanostructure

**DOI:** 10.1101/2024.09.02.610702

**Authors:** Felix Barber, Zarina Akbary, Zhe Yuan, Jacob Biboy, Waldemar Vollmer, Enrique R. Rojas

## Abstract

The cell wall is a polymeric exoskeleton that defines the size and shape of bacteria; it is composed of peptidoglycan and, in Gram-positive bacteria, wall teichoic acid polymers. Two systems synthesize peptidoglycan in rod-shaped bacteria, which are found pervasively across bacterial clades: the multi-protein Rod complexes synthesize anisotropic peptidoglycan, which is required for rod shape because it reinforces the cell wall along its circumference^1–3^, whereas the non-essential enzyme PBP1 synthesizes isotropic peptidoglycan^1,4^ (Fig. 1A). In Gram-positive bacteria, rod shape also requires wall teichoic acids^5^ for unknown reasons. Here, we show that wall teichoic acids promote rod shape by preventing the formation of nanoscopic pores in the *Bacillus subtilis* cell wall, which lead to amorphous growth by activating PBP1 and inhibiting Rod complexes. Depleting wall teichoic acids resulted in pores within minutes, coinciding with a rapid increase in PBP1-mediated synthesis. PBP1’s ability to sustain teichoic acid-less growth depended on its intrinsically disordered domain. In contrast to previous steady-state measurements^6,7^, we found that wall teichoic acid depletion caused the transient arrest of Rod complexes prior to the onset of amorphous growth. Finally, one of the two synthetically lethal cell wall hydrolases in *B. subtilis*^8^, LytE, became essential during wall teichoic acid depletion, meaning that PBP1 and LytE execute a novel, amorphous mode of growth. Collectively, our results identify the cell wall, via its molecular-scale structure, as a non-canonical auto-regulator of its own synthesis.

**Fig. 1:**
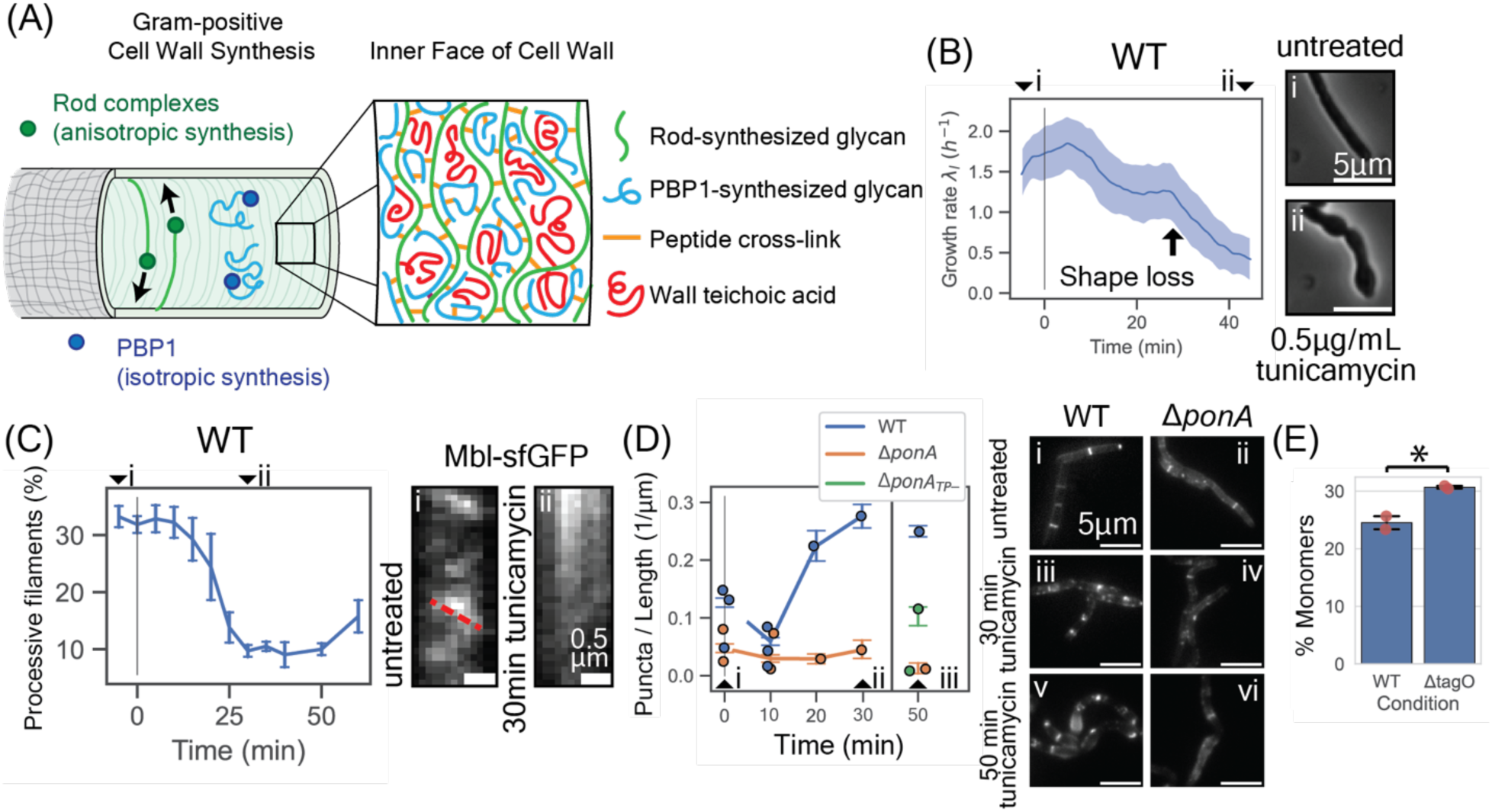
Inhibiting wall teichoic acid synthesis decreases Rod complex activity prior to cell shape loss. (A) Schematic of Gram-positive cell wall synthesis. **(B)** Cell length growth rate 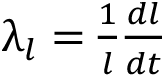 for wild-type cells during 0.5 𝜇g/mL tunicamycin treatment. Blue line shows smoothed population median, error bars show standard deviation. Data shown for 2,048 discrete cell tracks from 3 biological replicates. Inset: micrographs taken (i) before and (ii) after 45 min tunicamycin treatment. Scale bars 5 𝜇m. **(C)** Time course for the percentage of processive Mbl filaments during 0.5 𝜇g/mL tunicamycin treatment. Error bars show 95% confidence intervals across biological replicates (bootstrap analysis). Analysis performed on 62,901 discrete filament tracks from 4 biological replicates. Inset shows representative fluorescent kymographs of Mbl motion across the cell waist in (i) LB and (ii) after 30 min tunicamycin treatment. Red dotted line follows a processive Mbl filament. Scale bars 1 𝜇m. **(D)** Mean fluorescent puncta per unit cell length for wild-type, Δ*ponA* and *ponA_TP-_*cells during tunicamycin treatment, labeled with fluorescent D-amino acids. Error bars show 95% confidence intervals. Analysis performed over 2,017 segmentations from 4 biological replicates (wild-type), 857 segmentations from 3 biological replicates (Δ*ponA*) and 257 cells from 2 biological replicates (*ponA_TP-_*). All timepoints show statistical significance for difference in mean between cell types (Student’s T-test, P<0.01). Dots show individual replicates. Measurements at 50 min tunicamycin treatment used a lower exposure time so are plotted separately. Inset shows micrographs of (i,iii,v) wild-type and (ii,iv,vi) Δ*ponA* cells at (i,ii) 0 min, (iii,iv) 30 min, and (v,vi) 50 min of treatment. Micrographs are identically saturated across cell types. Scale bars 5 𝜇m. **(E)** Percentage of peptidoglycan subunits in monomers, measured by HPLC (*Methods*). Two biological replicates per strain. Mean values calculated across biological replicates. Error bars show 95% confidence intervals. Individual datapoints shown in red. Statistical significance calculated using Student’s T-test, P<0.05.

## Introduction

The bacterial cell wall is a solid polymeric macromolecule that protects cells from lysis in many contexts, including osmotic shock^9^, pathogenesis^10^ and antibiotic treatment^11^. The cell wall also prescribes cell shape (Fig. 1A), which affects many aspects of cellular physiology^3,12,13^ — a pervasive shape across bacteria is the bacillus, or “rod”^14^. The cell wall consists primarily of peptidoglycan: glycan polymers that are covalently crosslinked through short peptides. In Gram-positive bacteria, peptidoglycan is also covalently modified with wall teichoic acids, anionic polymers of sugar alcohol-phosphate subunits (hereafter “teichoic acids,” distinct from the membrane-anchored lipoteichoic acids)^15,16^. Teichoic acids are usually nonessential^5,17^, however, inhibiting their synthesis causes *Bacillus subtilis* and *Listeria monocytogenes* to grow slowly^5^ and amorphously^17,18^. How teichoic acids enable rod-shaped morphogenesis is unknown^5,17,18^.

During rod-shaped cell growth, peptidoglycan is synthesized by two molecular mechanisms. First, the essential, multi-protein Rod complexes synthesize anisotropic peptidoglycan that structurally reinforces the cell wall along its circumference (Fig. 1A), providing the mechanical basis for rod shape^1,2^. Anisotropic peptidoglycan synthesis results from the processive circumferential motion of Rod complexes along the plasma membrane^19–21^. In *B. subtilis*, this motion is oriented by actin homologs MreB, Mbl and MreBH^22,23,6^, and is driven by peptidoglycan synthesis itself^19^. Circumferential motion, therefore, is a direct indicator for Rod complex activity^24,25^. Within the Rod complex, peptidoglycan polymerization is catalyzed by the transglycosylase RodA^26^, drawing from lipid II precursors in the plasma membrane, while cross-linking of nascent polymers to acceptor peptides within the existing cell wall is catalyzed by the transpeptidases PBP2A and PBPH^4,27^ (“Class B” PBPs).

The second mechanism is the non-essential, multi-domain enzyme PBP1 (a “Class A” PBP, encoded by *ponA*^28^) that synthesizes isotropic peptidoglycan (Fig. 1A). PBP1 is thought to repair pores in the cell wall that arise during the rapid cell-wall turnover required for cell growth^4,29^. Because Rod complexes synthesize peptidoglycan anisotropically and PBP1 synthesizes it isotropically, the balance of their expression determines cell width^1^, and overexpression of PBP1 causes semi-amorphous growth^1^.

Teichoic acids influence diverse cellular phenotypes including antimicrobial resistance^30,31^, protein localization within the cell envelope^32^, pathogenesis^33^, cation homeostasis^34,35^ and autolysis^36–39^. It is unclear, however, how these processes would account for the dependence of cell shape on teichoic acids according to the Rod complex-PBP1 model. One possibility is that Rod complexes are directly, biochemically activated by teichoic acids^40^. This model is consistent with the observations that teichoic acid synthases and peptidoglycan synthesis machinery both exhibit punctate spatial patterning^41^, and that the syntheses of the two materials are co-local in both *B. subtilis*^42^ and *Streptococcus pneumoniae*^43^. However, teichoic acid synthases do not exhibit the circumferential motion characteristic of Rod complexes, and Rod complexes move processively along the cell membrane in amorphous cells depleted for TagO, the enzyme that catalyzes the first committed reaction of teichoic acid synthesis^6,7^. Given these conflicting data, there is no known mechanistic interaction between teichoic acids and either of the two peptidoglycan biosynthesis pathways.

## Results

### Inhibiting wall teichoic acid synthesis reduces growth rate prior to cell shape loss

*Bacillus subtilis* mutants that lack teichoic acids (Δ*tagO*) grow amorphously and ≈4 times slower than wild-type cells in bulk liquid culture^5^. To interrogate this dependence at the single-cell level, we measured the dynamics of growth rate and cell shape upon teichoic acid depletion by using microfluidics to acutely treat exponentially growing, wild-type *B. subtilis* cells with the small molecule tunicamycin (Fig. 1B), which specifically inhibits TagO at low concentrations^31,44–46^ (Fig. S1A,B,C). Approximately 10 minutes after tunicamycin treatment, the median growth rate decreased from its steady-state value in rich medium (LB), plateaued temporarily, and then decreased a second time until it reached its steady-state value in tunicamycin (Fig. 1B, S2A). Cells retained their untreated rod shape until the second decrease in elongation rate when they began to widen, eventually losing rod shape altogether (Fig. S2B). Amorphous cells grew indefinitely (Video S1). We observed a similar decline in the single-cell growth rate prior to the loss of cell shape when we acutely inhibited *tagO* transcription (Fig. S2C,D), during tunicamycin treatment on agarose pads (Fig. S2E,F), and during tunicamycin treatment in minimal media (S750; Fig. S2G,H; *Methods*).

### Inhibiting wall teichoic acid synthesis reduces Rod complex activity

Although it was previously reported that inhibiting teichoic acid synthesis does not prevent Rod complex motion during steady-state, amorphous cell growth^6,7^, we hypothesized that the loss of cell shape results from a quantitative or transient effect of teichoic acid depletion on Rod complex activity. To test this, we measured the dynamics of Rod complex motion upon tunicamycin treatment by using total internal reflection fluorescence microscopy to track a fusion of superfolder GFP and Mbl^19,24,25^, a protein scaffold of the Rod complex. As we hypothesized, tunicamycin caused an acute decrease in the percentage of Rod complexes that moved processively, (Fig. 1C), in the spatial density of processive Rod complexes (Fig. S3A), and in Rod complex speed (Fig. S3B). These decreases were coincident with the first decline in cellular growth rate (Fig. 1B). Following the second decrease in growth rate (and loss of cell shape; Fig. 1B) we observed a small but reproducible recovery of Rod complex motion (Fig. 1C) against a higher background of diffuse Mbl-sfGFP fluorescence than we observed in untreated cells (Fig. S3C, Videos S2-S7), consistent with the previous observation of Rod complex motion in Δ*tagO* mutants^6^. Tunicamycin also inhibited Rod complex motion as tracked via another Rod complex scaffold (MreB-mNeonGreen; Fig. S3D-F), and inhibited Rod complexes in slow-growing cells in minimal media (S3G-I; *Methods*). Finally, like tunicamycin treatment, transcriptional inhibition of *tagO* expression caused a decrease in each metric for Rod complex activity (Fig. S3J-L). The decrease was more gradual for transcriptional inhibition, which we expected since in this case TagO enzymes must be diluted by growth before teichoic acids are depleted.

Since Rod complexes synthesize peptidoglycan uniformly across the cell length, we hypothesized that this distributed synthesis would be perturbed upon the Rod complex arrest associated with teichoic acid depletion. To test this, we measured the localization of peptidoglycan synthase activity. First, we pulsed exponentially growing cells with the fluorescent D-amino acid 7-hydroxycoumarin-3-carboxylic acid-amino-D-alanine (HADA), which is incorporated into peptidoglycan by transpeptidases^47,48^, for 5 min at various times after tunicamycin treatment. Whereas in untreated cells HADA was incorporated uniformly across the cell wall, tunicamycin treatment caused HADA to be incorporated as aberrant puncta (Fig. 1D), consistent with a loss of distributed Rod complex synthesis. Tunicamycin had the same effect on the dipeptide D-amino acid ethynyl-D-alanine D-alanine (EDA-DA), which is incorporated into peptidoglycan precursors cytosolically^49^ (Fig. S4B). Finally, transcriptional inhibition of *tagO* also caused punctal HADA incorporation (Fig. S4A).

Since Rod complexes are the primary mode of peptidoglycan synthesis during vegetative growth, we hypothesized that a loss of Rod complex activity during teichoic acid depletion would require increased peptidoglycan synthesis by another mechanism, and that this shift would cause changes in the abundance of specific types of peptidoglycan crosslinks. To test this, we performed high performance liquid chromatography on purified, lysozyme-digested peptidoglycan from exponentially growing wild-type and Δ*tagO* cells^50^. Deleting *tagO* caused a small increase in the percentage of un-cross-linked peptidoglycan subunits, and a corresponding decrease in the percentage of dimeric cross-links (Fig. 1E, S5A). Conversely, we observed increases in trimeric and tetrameric cross-links in Δ*tagO* cells that were reproducible but not significant across two replicates (Trimers: 9.9% ± 0.1% for Δ*tagO* vs. 8.0% ± 0.7% for wild-type. Tetramers: 1.6% ± 0.2% for Δ*tagO* vs. 0.8% ± 0.1% for wild-type. Error is s.e.m.; Fig. S5B,C,D). The correlation of trimeric and tetrameric cross-linking with reduced Rod complex activity is consistent with a previous study that reported the reduction of these species in Δ*ponA* cells^50^.

### PBP1 is essential in the absence of wall teichoic acids

Since i) Rod complexes and PBP1 are the two principal modes of peptidoglycan synthesis during cell growth, ii) teichoic acid depletion acutely inhibits Rod complex activity (Fig. 1C), and iii) cells lacking teichoic acids grow amorphously with a cell wall that is rich in trimeric and tetrameric cross-links, as would be expected from PBP1-mediated cell growth, we hypothesized that teichoic acid-less growth is driven by PBP1. To test this, we first measured the effect of tunicamycin treatment on the growth, shape, and viability of Δ*ponA* mutants. At the single-cell level, tunicamycin caused a decrease in the growth rate of Δ*ponA* cells equivalent to the growth-rate decrease it caused in wild-type cells, except without the transient plateau and subsequent shape loss (Fig. 2A). Rather, Δ*ponA* mutants exhibited pervasive lysis after their growth rate decrease (Fig. 2B). Furthermore, concentrations of tunicamycin that permitted amorphous growth of wild-type cells prevented growth of Δ*ponA* cells (Fig. 2C, S6A).

**Figure 2:**
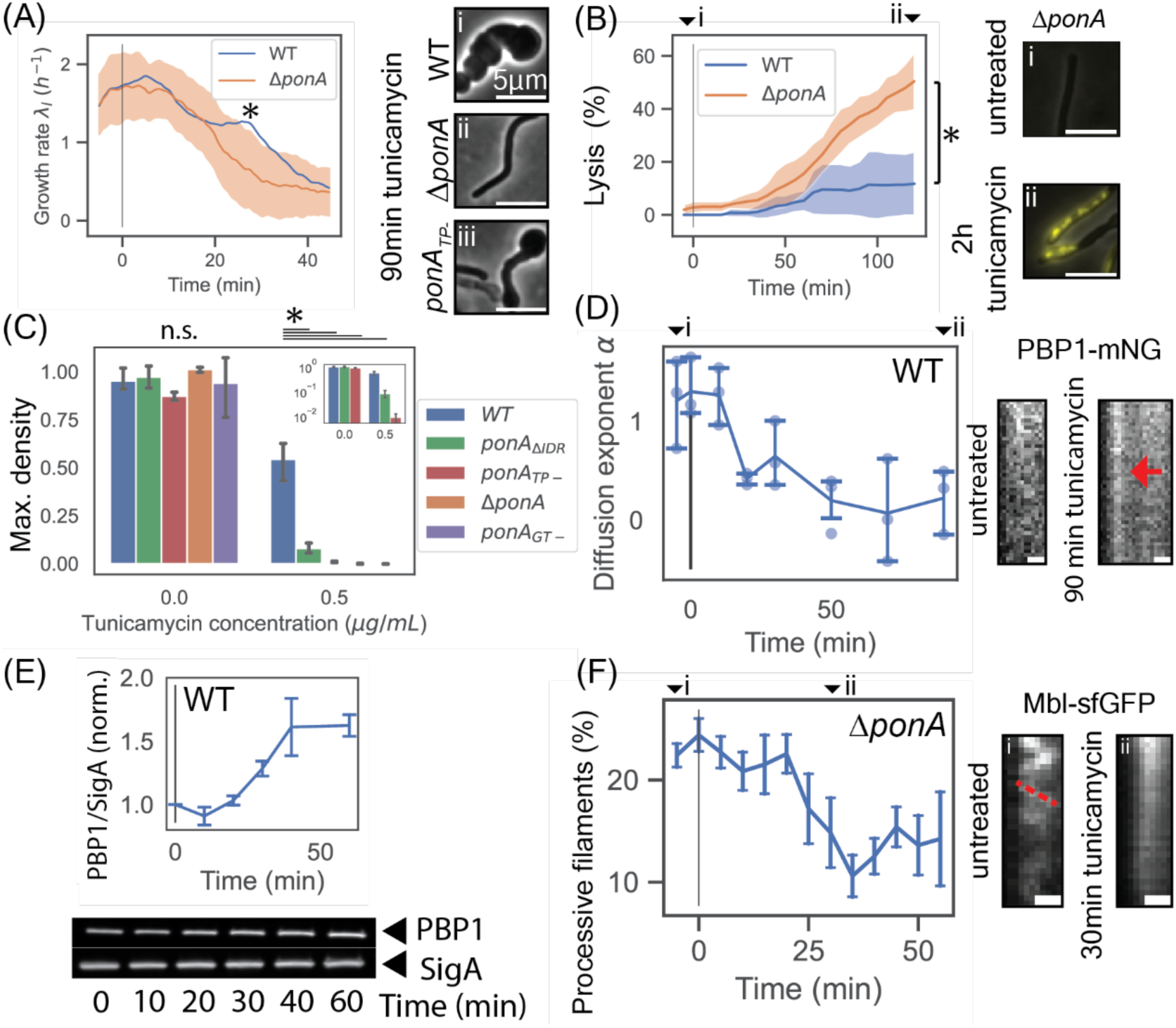
P**B**P1 **is required for growth during wall teichoic acid synthesis inhibition. (A)** Cell length growth rate 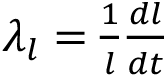 for Δ*ponA* cells during 0.5 µg/mL tunicamycin treatment. Orange line shows smoothed population median, error bars show standard deviation. Analysis performed over 629 discrete cell tracks from 5 biological replicates. Blue line reproduces wild-type data from Fig. 1A. Statistical significance calculated between cell types at 25 min tunicamycin exposure using Student’s T-test, P<0.01. Inset: micrographs showing (i) wild-type, (ii) Δ*ponA* and (iii) *ponA_TP-_* cells following 90 min treatment with 0.5 µg/mL tunicamycin. Scale bars 5 µm. **(B)** Percentage of membrane-disrupted cells following tunicamycin exposure, measured by propidium iodide staining. Solid line shows mean across all fields of view. Error bars show standard deviation of the same. Analysis performed over 1,098 cells from three biological replicates (Δ*ponA*) and 1,105 cells from two biological replicates (wild-type). Statistical significance calculated for final timepoint using Student’s T-test, P<0.01. Inset: micrographs of *ΔponA* cells (i) before and (ii) after 2 h tunicamycin treatment. Phase images overlaid with propidium iodide stain. Scale bars 5𝜇m. **(C)** Maximum OD (optical density) for wild-type, *ponA*_Δ*IDR*_ (expressing PBP1 without its intrinsically disordered region)*, ponA_TP-_* (expressing PBP1 with an inactive transpeptidase domain), Δ*ponA* and *ponA_GT-_*(inducibly expressing PBP1 with an inactive glycosyltransferase domain) with and without a low dose of tunicamycin, grown in bulk culture. Data shown from one representative biological replicate out of three, with four technical replicates per biological replicate. Error bars show 95% confidence intervals based on bootstrap analysis. Inset: same data for WT, *ponA*_Δ*IDR*_ and *ponA_TP-_* cells, plotted on log-scale. Statistical significance tested with one-way ANOVA followed by Tukey’s HSD post-hoc test for groups with and without tunicamycin, P<0.01. **(D)** Diffusion exponent 𝛼 for PBP1 puncta during a time course of tunicamycin treatment (*Methods*). 𝛼 = 1: diffusive, 𝛼 < 1: sub-diffusive. Inset: kymographs of PBP1-mNeonGreen motion for (i) untreated and (ii) 90 min tunicamycin-treated cells. Red arrow points to a static PBP1-mNeonGreen spot during tunicamycin treatment. Total time: 4 s. **(E)** Western blot analysis of PBP1 levels during 0.5µg/mL tunicamycin treatment. Bottom: Western blot bands for PBP1-FLAG and identically loaded SigA control. Loading volumes normalized based on OD600 readings. Top: Quantification of PBP1 staining, normalized by SigA stain and measured relative to values at 0min tunicamycin treatment. Two biological replicates. Error bars show standard error of the mean. **(F)** Percentage of processive Mbl-sfGFP filaments during tunicamycin time course of Δ*ponA* cells. Error bars show 95% confidence intervals across biological replicates based on bootstrap analysis. Analysis performed on 40,793 discrete filament tracks from three biological replicates.

Teichoic acid-less cell growth was dependent on the glycosyltransferase activity of PBP1, since a mutant incapable of this reaction (*ponA_GT-_*) did not grow at concentrations of tunicamycin that permitted wild-type growth (Fig. 2C; *Methods*). Similarly, tunicamycin inhibited the growth of a PBP1 mutant incapable of transpeptidation (*ponA_TP-_*; Fig. S6B)^51^ 40-times more strongly than wild-type growth (Fig. 2C), and caused only partial shape loss of this mutant prior to growth arrest (Fig. 2A inset). Tunicamycin may inhibit the growth of the *ponA_GT-_* mutant more than it inhibits that of the *ponA_TP-_* mutant because PBP1 transpeptidase activity is dependent on glycosyltransferase activity^52^. In addition to its catalytic domains, PBP1 also possesses a highly anionic 14kDa C-terminal intrinsically disordered domain, which stimulates PBP1-mediated synthesis upon cell wall damage^29^; we found that tunicamycin inhibited the growth of a mutant harboring a version of PBP1 lacking this domain (*ponA*_Δ*IDR*_; Fig. 2C, S6B). Finally, transcriptional inhibition of *tagO* expression in Δ*ponA* and *ponA_TP-_* mutant backgrounds prevented growth in bulk culture (Fig. S6C).

These results demonstrate that PBP1 activity is essential for the amorphous cell growth following teichoic acid depletion, which inhibits Rod complexes. In this light, we hypothesized that amorphous growth would require increased peptidoglycan synthesis by PBP1 upon teichoic acid depletion, concomitant with the decline in Rod complex activity (Fig. 1C). To test this, we performed single-molecule tracking on a PBP1-mNeonGreen fusion protein expressed at low levels, in addition to the native enzyme (Fig. 2D, S6D; *Methods*). Previous studies demonstrated that PBP1 is stationary when it is actively synthesizing peptidoglycan and rapidly diffuses when it is not^4^. Consistent with our hypothesis, we observed a stark decline in PBP1 diffusion upon tunicamycin treatment, culminating in large, stable PBP1 puncta (Fig. 2D, Videos S8-9).

To confirm that these PBP1 puncta were enzymatically active, we performed HADA staining after 90 minutes of tunicamycin treatment and measured the colocalization of PBP1 and HADA puncta. We found that 67% of PBP1 puncta overlapped with HADA puncta, while a simulated random localization yielded an overlap of only 36% (Fig. S6E). Furthermore, Δ*ponA* cells treated with tunicamycin did not display HADA puncta (Fig. 1D), indicating that these puncta resulted from PBP1-polymerized peptidoglycan. However, *ponA_TP_*_-_ cells exhibited only a partial reduction in HADA puncta labelling compared to wild-type (Fig. 1D), suggesting that other transpeptidases may partially complement PBP1 transpeptidation during teichoic acid depletion. This is consistent with the incomplete growth inhibition of tunicamycin-treated *ponA_TP-_*cells, as opposed to the full arrest of tunicamycin-treated Δ*ponA* and *ponA_GT-_* cells (Fig. 2C).

To test whether PBP1 expression increases in concert with PBP1-mediated synthesis upon teichoic acid depletion, we measured PBP1 abundance by western blot (Fig. 2E), and by labeling with the fluorescent penicillin bocillin, which covalently binds transpeptidases at their active sites^53^ (Fig. S6F-H). Upon tunicamycin treatment, we observed a similar increase in PBP1 levels as measured with each assay (62% ± 8% by western blot, 80% ± 6% by bocillin labeling. Errors are s.e.m.). Notably, the increase in PBP1 bocillin labeling preceded the increase in PBP1 staining by western blot (Fig. S6H), on the same timescale as decreased PBP1 diffusion/synthesis. The increase in PBP1 expression did not result from the SigI stress response that is triggered by teichoic acid depletion^54^, since we also observed increased PBP1 expression in Δ*sigI* mutant cells (Fig. S6I).

Our results demonstrate that teichoic acid depletion causes an increase in PBP1 abundance and PBP1-mediated synthesis, and that this enzyme becomes essential for growth without teichoic acids. Based on previous studies^1^, the increase in PBP1 expression that we measured is, alone, insufficient to cause the degree of cell shape changes that we observed. Since our single-molecule measurements indicate that PBP1-mediated synthesis increases upon teichoic acid depletion, our data are consistent with a model in which teichoic acids inhibit peptidoglycan synthesis by PBP1 such that when teichoic acids are depleted PBP1 outcompetes Rod complex for lipid II precursors leading to slow, amorphous cell growth and Rod complex inhibition.

Given this model, we next questioned whether Rod complex motion also decreases in the absence of PBP1. We assayed Rod complex motion during tunicamycin treatment in Δ*ponA* cells (*Methods*), finding that Rod complex motion declined, but with a delay compared to wild-type cells (Fig. 2F, Videos S10-S11). In this case, a mechanism besides competition for precursors must inhibit Rod complex function during teichoic acid depletion.

### Teichoic acid depletion results in a porous cell wall in the absence of PBP1

We next asked why PBP1 becomes activated during teichoic acid depletion, and why Rod complexes arrest during teichoic acid depletion in Δ*ponA* cells. Since PBP1 is thought to fill pores in the cell wall^4^ and is conditionally essential in the absence of teichoic acids, we hypothesized that teichoic acids play a complementary pore-filling role and that Rod complexes require a pore-less “confluent” cell wall on which to synthesize new peptidoglycan. Furthermore, since intrinsically disordered domains sense cell wall damage^29^, likely through entropic translocation through pores in the cell wall^55^, and deleting PBP1’s intrinsically disordered region prevents growth during teichoic acid depletion (Fig. 2C), we additionally hypothesized that PBP1-mediated synthesis is stimulated by this domain when it senses natural pores in the teichoic acid-less cell wall.

To test whether depleting teichoic acids results in pores within the cell wall, we leveraged a single-cell microfluidics assay we recently developed^56^. In this assay, we used a brief pulse of detergent to lyse cells expressing genetically encoded, cytosolic fluorescent probes and then measured the kinetics of the probes’ diffusion through the cell wall at the single-cell level — this provides an empirical measurement of cell wall permeability (*Methods*). To measure the pore structure at different length scales, we used two probes: mNeonGreen (27.5 kDa, minimal radius *R*_min_≈2.0 nm^57^) and a new fluorogenic version of ubiquitin that we developed (ubiquitin-FlAsH, 9.5 kDa, *R*_min_≈1.4 nm).

We first measured the effect of teichoic acid depletion on cell wall permeability by treating wild-type and 11*ponA* cells with tunicamycin (Fig. 3A). To avoid confounding effects on cell wall permeability caused by changes in peptidoglycan synthesis, we measured permeability before Rod complex activity began to decline (10 min for WT and 20 min for 11*ponA*). Contrary to our initial hypothesis, tunicamycin treatment of wild-type cells resulted in a modest but reproducible decrease in the permeability of wild-type cell walls, as measured with the mNeonGreen probe (Fig. 3A, S7A). Conversely, the same treatment resulted in an increase in permeability in 11*ponA* cells.

**Figure 3:**
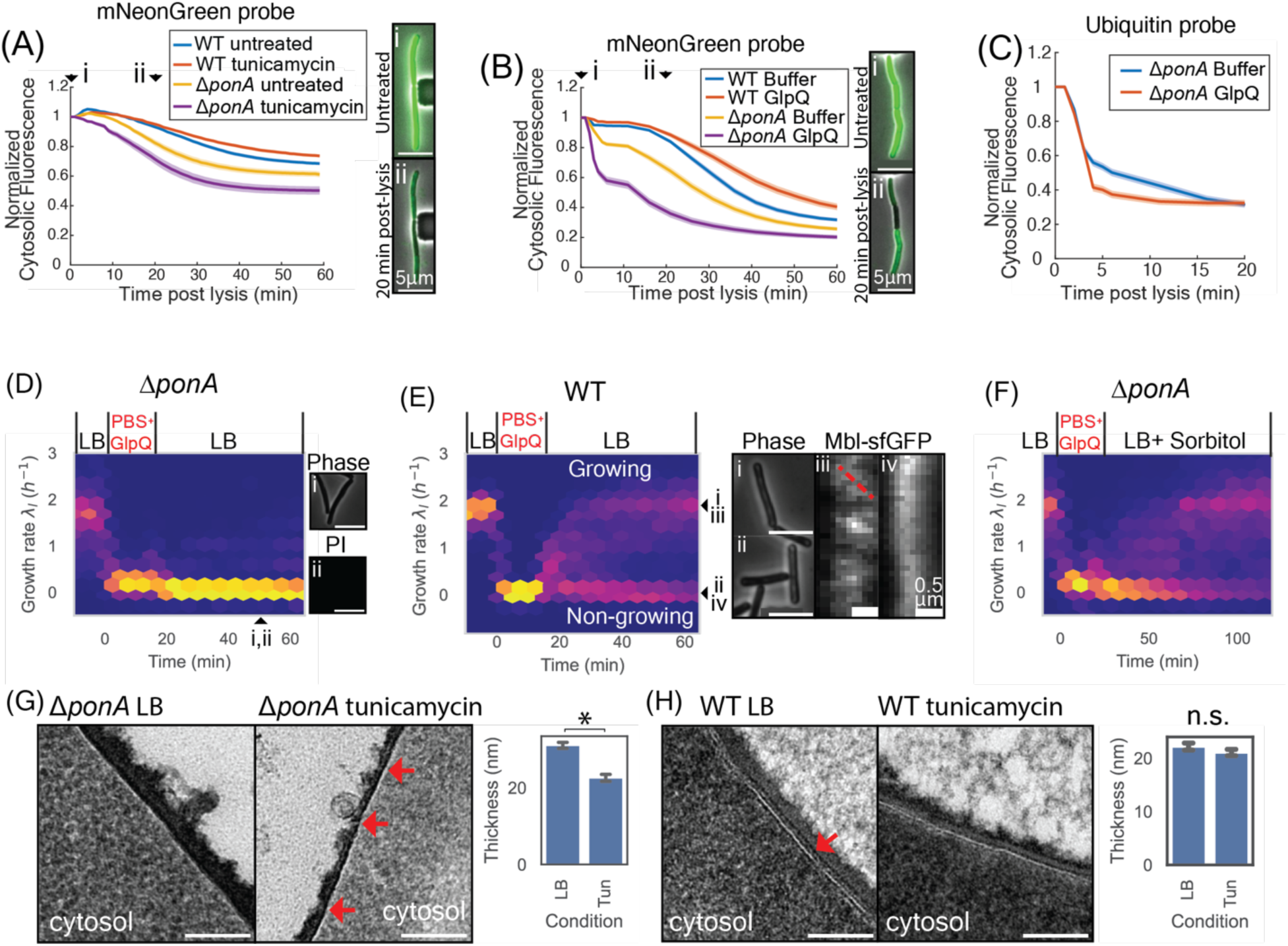
PBP1 fills pores that are exposed upon wall teichoic acid removal, conferring growth. **(A)** mNeonGreen diffusion out of cells with disrupted membranes following a 10 min (wild-type) or 20 min (Δ*ponA*) incubation with LB supplemented with tunicamycin or with LB alone. Solid lines show population average, error bars show standard error of the mean. Wild-type untreated: 87 cells, 4 biological replicates. Wild-type tunicamycin: 139 cells, 3 biological replicates. Δ*ponA* untreated: 53 cells, 3 biological replicates. Δ*ponA* tunicamycin: 35 cells, 3 biological replicates. Inset: micrographs showing cellular fluorescence of tunicamycin-treated Δ*ponA* cells (i) before and (ii) 20 min after cell lysis. Scale bars 5 𝜇m. **(B)** mNeonGreen diffusion out of lysed wild-type and Δ*ponA* cells following a 15-minute live-cell incubation with PBS supplemented with either 40 µM GlpQ in Buffer A or with Buffer A alone. Wild-type Buffer: 121 cells, 1 biological replicate. Wild-type GlpQ: 90 cells, 1 biological replicate. Δ*ponA* Buffer: 85 cells, 3 biological replicates. Δ*ponA* GlpQ: 108 cells, 3 biological replicates. Inset: micrographs showing cellular fluorescence of GlpQ-treated Δ*ponA* cells (i) before and (ii) 20 min after cell lysis. Scale bars 5𝜇m. **(C)** Ubiquitin-FlAsH diffusion out of lysed Δ*ponA* cells following a 2 min incubation with PBS + 5% N-lauroylsarcosine supplemented with either 40 µM GlpQ or the equivalent volume of Buffer A. **(D)** Heat map showing growth rate 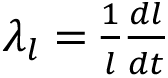 of Δ*ponA* cells following a 15 min incubation with 10µM native GlpQ (1,816 discrete tracks, 3 biological replicates). Heat map intensity is proportional to the number of cells with a given growth rate at a given time, normalized across the total number of cells tracked at each timepoint. Inset: lack of propidium iodide staining for Δ*ponA* cells 50 min after GlpQ incubation. (i) Phase contrast. (ii) Propidium iodide stain. For positive stain, see Fig. 2B ii. Scale bars 5µm. **(E)** Same as (D) for wild-type cells (3,236 discrete tracks, four biological relicates). Inset: (i-ii) cell micrographs of (i) growing and (ii) non-growing cells, and (iii-iv) kymographs of (iii) growing and (iv) non-growing cells, following the exit from GlpQ treatment. Scale bars 5𝜇m in (i,ii), 0.5 𝜇m in (iii,iv). Red dotted line in (iii) follows a processive Mbl filament. **(F)** Heatmap showing Δ*ponA* cell length growth rate 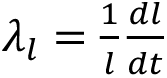 when exit from a 15-minute incubation with 10 µM GlpQ was coupled to a 500 mM hyperosmotic shock (3,036 discrete cell tracks, 3 biological replicates). **(G-H)** Transmission electron micrographs showing the sidewall of exponentially growing cells grown in either LB or LB supplemented with 0.5 𝜇g/mL tunicamycin for 30 min. Scale bars 100 nm. **(G)** Δ*ponA* cells. Red arrows in show sites of severe cell wall thinning during tunicamycin treatment. Inset: average wall thickness for the growth conditions shown, calculated for 47 cells (LB) and 44 cells (tunicamycin). Statistical significance calculated using Student’s T-test, P<0.01. **(H)** Wild-type cells. Red arrow shows electron-dense band on the inner face of the cell wall, absent in tunicamycin-treated cells. Inset: average wall thickness of wild-type cells for the growth conditions shown, calculated for 31 cells (LB) and 28 cells (tunicamycin). No statistically significant difference measured between conditions, Student’s T-test for P<0.01.

As an alternative approach to remove teichoic acids, we perfused cells with a purified phosphodiesterase, GlpQ, that *B. subtilis* naturally expresses during phosphate starvation to digest and recycle its teichoic acids^58^ (*Methods*). Perfusing cells with GlpQ dissolved in phosphate buffer saline (PBS) for 15 minutes reduced the abundance of teichoic acids within the cell wall (Fig. S8; *Methods*) and recapitulated the effect of tunicamycin on cell wall permeability: it caused a modest decrease in mNeonGreen permeability in wild-type cells but a large increase in permeability in 11*ponA* cells (Fig. 3B, S7B). Together, these results are consistent with a model in which teichoic acids occlude pores in the cell wall, and in which PBP1 is activated by these pores via its intrinsically disordered domain to “seal” them, thereby preventing an increase in cell wall permeability^56^.

Treating 11*ponA* cells with tunicamycin or GlpQ for durations approaching the cell-cycle time (≈20 min) could cause pore formation indirectly, for example, by inducing peptidoglycan hydrolysis unbalanced by synthesis^36–39^. To control for this possibility, we measured cell-wall permeability during teichoic acid digestion by GlpQ, reasoning that at short time scales after teichoic acid digestion, peptidoglycan hydrolases would not have had time to meaningfully enlarge pores. Because we anticipated small, incipient pores, we used our ubiquitin probe to test for them. When we simultaneously digested teichoic acids and measured the loss of this probe in Δ*ponA* cells, we observed an increase in cell wall permeability within 2 minutes of GlpQ treatment (Fig. 3C, S7C, S9A,B).

In concurrent work, we discovered that the size threshold for cell wall permeability is close to the size of typical globular proteins^56^. We therefore hypothesized that the pores exposed during teichoic acid depletion would cause increased labeling by peptidoglycan-binding proteins. To test this, we measured whether teichoic acid depletion affected cell-wall labeling with a fluorescent lectin that specifically binds to the N-acetyl glucosamine moiety of peptidoglycan. We observed an increase in labeling of both Δ*ponA* cells and, to a lesser extent, wild-type cells following pre-treatment with tunicamycin (Fig. S9C). We speculate that teichoic acid depletion in wild-type cells reduces cell wall permeability (Fig. 3A) but increases lectin binding because PBP1 is membrane-bound and would therefore only fill pores near the internal surface of the cell wall, leaving teichoic acid-less peptidoglycan on the outer surface exposed.

Our model that Rod complexes require a pore-less cell wall as their substrate made the prediction that removing teichoic acids from 11*ponA* cells by perfusing them with PBS-suspended GlpQ, which arrests cell growth and leads to a permeable cell wall, would prevent Rod complex activity and cell growth upon re-immersion in growth media. Consistent with this prediction, 11*ponA* cells were unable to recover from teichoic acid digestion but did not lyse upon re-immersion in growth media (Fig. 3D), whereas most wild-type cells recovered (Δ*ponA*: 7% ± 2% recovered. Wild-type: 66% ± 2% recovered. Recovery counted as reaching half the initial, unperturbed growth rate. Error is s.e.m.; Fig. 3E). Furthermore, wild-type cells that resumed growth did so as rods and recovered Rod complex activity (Fig. 3Ei,iii), while cells that did not recover showed no Rod complex motion (Fig. 3Eii,iv, S10A-D). Rod complex activity and cell growth of both wild-type and Δ*ponA* cells recovered fully from incubation in PBS alone (Wild-type: 94% ± 1% recovered growth. Δ*ponA*: 81% ± 2% recovered growth. Error is s.e.m.; Fig. S10E-G, S11A,B). This correlation between pores, Rod complex activity and PBP1 is strong evidence that the Rod complexes cannot add peptidoglycan to a porous cell wall. However, since GlpQ-treatment of wild-type cells does not induce pores (Fig. 3C) but partially inhibits growth recovery (Fig. 3E), teichoic acid digestion must additionally inhibit Rod complexes via a mechanism independent of pores, at least of those that are detected by the mNeonGreen probe. We speculate that this results from the effect of spatially heterogeneous peptidoglycan synthesis by PBP1 on Rod complexes (*Discussion*).

We reasoned that if pores in the cell wall inhibit Rod complex activity upon teichoic acid removal, then shrinking these pores would activate Rod complexes, thereby promoting cell growth recovery after GlpQ treatment. We tested this prediction by coupling the re-immersion of cells in rich media following GlpQ treatment to a 500 mM hyperosmotic shock that physically contracted the cell wall^59^. To ensure that any cell growth recovery resulted directly from cell wall contraction and not from PBP1 activity, we performed this experiment in Δ*ponA* cells that are otherwise unable to grow following teichoic acid cleavage. Consistent with our hypothesis, we found that hyperosmotic shock led to a partial growth recovery after teichoic acid removal even in the absence of PBP1 (60% ± 2% recovered. Error is s.e.m.; Fig. 3F, S11C).

Finally, we sought to visualize the effect of teichoic acid depletion on the cell wall. Previously, atomic force microscopy explicitly aimed at resolving peptidoglycan structure did not resolve a difference between isolated cell wall sacculi before and after hydrofluoric acid treatment to remove teichoic acids^60^. Therefore, we used transmission electron microscopy to examine the effect of teichoic acid depletion on cell wall ultrastructure. To do this, we performed high pressure freeze-substitution staining on chemically fixed wild-type and Δ*ponA* cells that had been grown either in LB alone or that had been treated for 30 min with tunicamycin (*Methods*). We observed significant cell wall degradation of Δ*ponA* tunicamycin-treated cell walls, visible both as general thinning of the cell wall relative to untreated cells, and as localized regions of lower density within the wall (Fig. 3G, S12A). Neither phenomenon was present in wild-type cells treated with tunicamycin (Fig. 3H, S12B). We also observed an electron-dense band on the inner face of the untreated cell wall that is absent in tunicamycin-treated cells (Fig. 3H), consistent with prior observations of a granular, teichoic acid-dependent layer supporting the Gram-positive periplasm in *Streptococcus pneumoniae*^61^. Although this technique is incapable of resolving the nanoscale pores detected by our permeability assays, these findings are consistent with the conclusion that PBP1 fills pores within the cell wall during teichoic acid depletion, and that in Δ*ponA* cells, teichoic acid depletion causes significant cell wall degradation, leading to a loss of Rod complex activity.

### Rod complex inhibition and shape loss occur during teichoic acid depletion in hydrolase mutants

We finally questioned whether the cell wall degradation that occurs upon teichoic acid depletion in *11ponA* cells (Fig. 3A,B,G, S12A) depends on specific peptidoglycan hydrolases. Although *Bacillus subtilis* encodes up to 42 putative hydrolases, cell growth can be sustained by either of two synthetically lethal D,L endopeptidases: CwlO or LytE^8^. Furthermore, teichoic acid depletion significantly increases SigI-dependent *lytE* expression^54^. Therefore, we measured the growth rate and width dynamics of Δ*cwlO* and Δ*lytE* cells upon tunicamycin treatment, in the presence or absence of PBP1, and found that the initial phases of these dynamics were identical to those of the respective background strains possessing both hydrolases (Fig. S13A-D). Additionally, upon tunicamycin treatment, Rod complex activity decreased in both hydrolase mutants, as it did in wild-type cells (Fig. S13E-J). Strikingly, at long times after either tunicamycin treatment (Fig. S14A), or transcriptional inhibition of *tagO* (Fig. S14B), Δ*lytE* cells failed to grow. This means that *lytE,* like *ponA,* is essential during teichoic acid depletion.

## Discussion

Our data support a model in which teichoic acids occlude nanoscopic pores in peptidoglycan (Fig. 4A,B) and prevent them from growing, which promotes rod shape by inhibiting PBP1 and enabling Rod complex activity. According to this model, in wild-type cells teichoic acids limit PBP1’s access to natural pores within peptidoglycan by “filling” them and/or by “lining” the inner face of the cell wall (Fig. 4C). When teichoic acids are depleted, PBP1’s intrinsically disordered domain rapidly senses the pores, leading to an increase in the relative amount of peptidoglycan that PBP1 synthesizes (Fig. 2D); we find it likely that the anionic teichoic acids exclude the anionic intrinsically disordered domain from pores in the cell wall through polymer-polymer repulsion, similar to the forces that govern macromolecular crowding^62^. Teichoic acid depletion also leads to an increase in PBP1 abundance (Fig. S6F-I) via an unknown mechanism. PBP1 is therefore the essential peptidoglycan synthase driving amorphous growth. This amorphous growth manifests gradually, via an intermediate “dumbbell” morphology that may reflect local reinforcement of the cell wall by septa^5^ (Fig. S15). Our model explains prior observations that Rod complexes become dispensable during teichoic acid depletion^63^.

**Figure 4:**
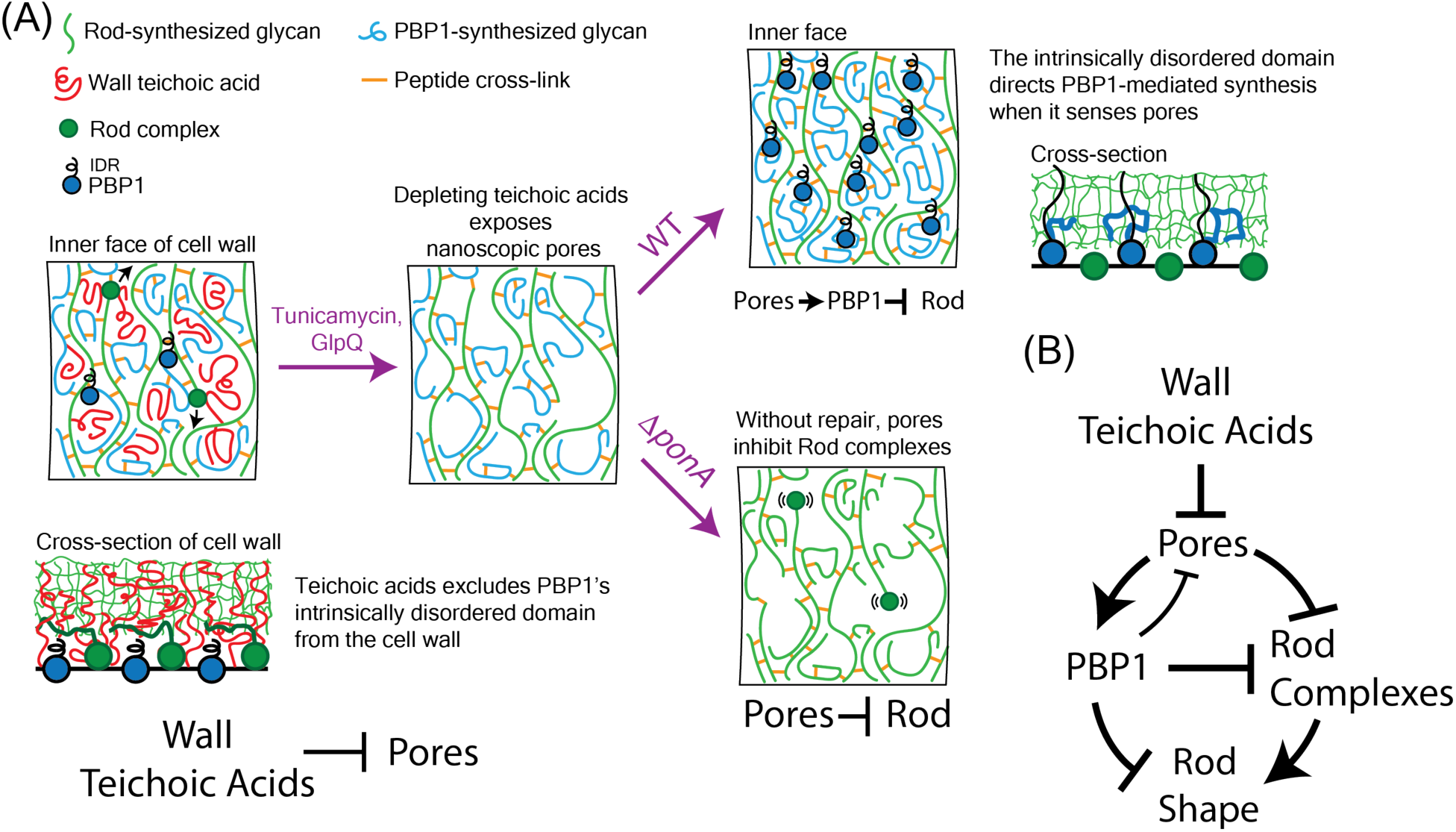
(A) Model illustration. During teichoic acid depletion of wild-type cells, denuded cell wall pores are targeted by PBP1, competitively inhibiting Rod complexes and driving localized peptidoglycan synthesis. This isotropic cell wall synthesis causes decreased growth and loss of cell shape. During teichoic acid depletion of Δ*ponA* cells, denuded cell wall pores directly arrest Rod complex motion through depletion of crosslinking acceptors. **(B)** Schematic of model described in (A).

At slightly longer times, teichoic acid depletion leads to decreased activity of Rod complexes (Fig. 1C). According to our model, this occurs in wild-type cells because increased PBP1-mediated synthesis inhibits Rod complex activity by reducing the lipid II pool. Reducing lipid II synthesis is known to inhibit Rod complex activity in both fast and slow growth conditions^25^, and we observed a reduction of Rod complex activity during teichoic acid depletion in both conditions (Fig. 1C, S3G-I). We speculate, however, that the heterogenous spatial distribution of PBP1-synthesized peptidoglycan upon teichoic acid depletion amplifies the inhibition of Rod complexes because one function of PBP1 in untreated cells is to contribute to a dense, homogeneous “lawn” of acceptor peptides. This hypothesis is consistent with the impaired Rod complex function of Δ*ponA* mutants during vegetative growth^64^, which we suppressed with supplemental magnesium (*Methods*). Localized PBP1 synthesis could inhibit Rod complexes if the latter could only cross-link efficiently to recently synthesized peptidoglycan (or peptidoglycan that is still being synthesized) due to either spatial or biochemical maturation of acceptor peptides (e.g. proximity to the plasma membrane). The heterogeneity of immature peptidoglycan would not inhibit PBP1 since it is not spatially processive, and can therefore cross-link to its own products.

In the absence of PBP1, teichoic acid depletion causes the simultaneous formation of nanoscopic pores in the cell wall (Fig. 3C) and the eventual arrest of Rod complexes (Fig. 2D,3D). According to our model, the pores exposed by teichoic acid depletion enlarge through hydrolysis without repair by PBP1 (Fig. 3A,B, 4A). When the pores become too large, the density of peptide acceptors becomes so low that Rod complexes arrest for the same reason that pharmacological inhibition of Class B PBP transpeptidation arrests Rod complexes^19–21^. Conversely, when these pores are contracted, Rod complex activity can recover (Fig. 3F). The delay in Rod complex arrest that we observed in 11*ponA* mutants compared to that in wild-type cells is consistent with competitive inhibition of Rod complexes by PBP1.

Our model accounts for why Rod complex activity partially recovers at late times after teichoic acid depletion, and in Δ*tagO* mutants. The transition from thin rods, to wide rods, to amorphous spheroids directly lowers the growth rate and the surface area-to-volume ratio of the cells, which is expected to increase lipid II levels by lowering the fraction of metabolic flux, per unit cell volume, needed for cell wall expansion. Rod complexes therefore become partially active again, but not at a high enough level to confer rod shape since PBP1 is constitutively de-inhibited and still directs peptidoglycan synthesis to nascent, teichoic acid-less pores.

In addition to cell wall synthesis, teichoic acids also influence peptidoglycan hydrolysis during cell growth, both directly and indirectly^36–39,54^. We found that *lytE* becomes essential during teichoic acid depletion for cells grown in LB, meaning that stable, amorphous teichoic acid-less growth is driven by PBP1 and LytE. However, deletion of *cwlO* or *lytE* had no effect on the growth, shape and Rod complex dynamics at early times after teichoic acid depletion, indicating that teichoic acids do not differentially regulate these two enzymes. Whether this conditional essentiality of *lytE* arises due to CwlO becoming inactive at long times following teichoic acid depletion will be the subject of further study. Our data suggests that in addition to directly occluding PBP1 from pores, teichoic acids also inhibit peptidoglycan hydrolysis during incubation in PBS (autolysis), potentially through inhibition of LytC and LytD^56^. Teichoic acids’ inhibition of hydrolysis could therefore underlie our observation that GlpQ-mediated digestion of teichoic acids partially prevents recovery of wild-type cells even though PBP1 activity prevents the formation of large pores (Fig. 3E). For example, runaway PBP1 activity and hydrolysis during GlpQ treatment could create an irreversibly altered cell wall incapable of supporting Rod complex activity.

*B. subtilis* cells depleted for wall teichoic acids often fail to separate and show aberrantly thick septa^5,6^.

Our data is consistent with increased PBP1-mediated synthesis contributing to septal peptidoglycan synthesis since Δ*ponA* cells show reduced septal HADA labeling relative to wild type cells during tunicamycin treatment (Fig. 1D). However, since teichoic acid-depleted cells that are forced to maintain rod shape through confinement grow and divide faster than their spherical counterparts^6^, the loss of rod shape clearly contributes substantially to this division defect.

In sum, wall teichoic acids are critical regulators of rod-shaped morphogenesis in Gram-positive bacteria since they coordinately influence both modes of peptidoglycan synthesis as well as its hydrolysis. Our study resolves the longstanding question of how teichoic acids promote rod-shape without directly contributing to cell wall anisotropy. Furthermore, we establish a direct link between the molecular structure of the cell wall and anisotropic cell wall synthesis, identifying the cell wall as a critical autoregulatory factor of its own synthesis, rather than a passive substrate.

## Methods

### Microscopy

All timelapse microscopy was performed with a Nikon Eclipse Ti2 microscope, outfitted with a photometrics PRIME BSI CMOS camera (Teledyne Photometrics) and a Nikon 100X Plan Apo Phase Contrast objective, NA 1.45. All non-TIRF fluorescence imaging used LED illumination. We conducted all imaging using CellASIC ONIX microfluidic plates (Millipore B04A-03), using standard priming and media exchange protocols. We heated all microscope stage components, CellASIC plates, and objectives to 37°C prior to imaging using a microscope incubator outfitted with a heater. We performed our timelapse phase contrast microscopy using 20s timesteps, monitoring medium exchange using the dye AlexaFluor 647 at a final media concentration of 2.5μg/mL, with CY5 illumination adjusted to avoid any confounding effects of phototoxicity.

### Total Internal Reflection Fluorescence (TIRF) Microscopy

All TIRF microscopy was performed with the same Nikon Eclipse Ti2 microscope, using a COHERENT 488nm laser and 100X Plan Apo objective, NA 1.45, with 2s timesteps. We adjusted our laser power, exposure time and TIRF alignment to minimize photobleaching, and used a fresh field of view for each new timepoint to avoid any confounding effects of phototoxicity.

### Single Molecule Tracking

We performed single molecule tracking by imaging small fields of view with TIRF microscopy, using a timestep of 90ms, laser power 1% and 40ms camera exposure time. To facilitate the rapid image acquisition, the laser remained on constantly for the 4s acquisition. To facilitate single-molecule resolution, we induced PBP1-mNeonGreen expression at low levels from a heterologous HyperSpank promoter with 10uM IPTG (Isopropyl ß-D-1-thiogalactopyranoside), alongside unperturbed native PBP1 expression from the endogenous locus. Note: Videos S8 and S9 were acquired with imaging conditions that visually emphasize the duration of PBP1 puncta during teichoic acid depletion, but that were not used for single-molecule tracking because of their low time-resolution: time step = 500ms, exposure time 50ms, 15s time course, 1% laser power. We generated our image file structures for downstream analysis using the ImageJ script tirf_pbp1_processing_v1.ijm. We then performed spot tracking using the python script single_molecule_tracking_v1.py, which detected and tracked filaments as follows: we detected puncta with SciKitImage’s bloblog feature, using manually curated thresholds to start each timelapse (that were then preserved within all timelapses acquired using that imaging condition). These thresholds were iteratively reduced throughout each step of a single time course to maintain roughly equivalent levels of spot detection. This allowed us to correct for partial photobleaching of the fluorescent signal. We tracked individual puncta across successive images (all analysed images used a timestep of 90ms) by calculating the minimum distance between all annotated puncta, subject to a maximum displacement threshold of two pixels (equivalent to a linear distance of 186nm based on our magnification and camera size). We then filtered to exclude all tracks with length 2 timepoints or less. Finally, we inferred the diffusion exponents by applying linear regression to log-log plots for the population-averaged mean-squared displacement of individual PBP1 puncta over time.

### Cell culture and transformation

Unless otherwise stated, all cell culture was performed with LB media (Luria Broth, Lennox). All *in vivo* imaging and spot-assay experiments were performed with log-phase cultures of OD600≤0.2. Transformations were performed with competent *B. subtilis* cells grown in MC Media + 20mM MgSO_4,_ then plated onto LB agarose plates. All cells were congenic with a PY79 strain background. Strains are tabulated in Table S1. We note that expression of *ponA_TG-_(E115A)* at high levels was toxic to cells, and required media supplementation with 30mM MgCl_2_. Similarly, our experiments on Rod complex tracking in Δ*ponA* cells were performed with 10mM supplemental MgCl_2_ to allow us to visualize processive Rod complexes in this strain background.

### S750 media

We used S750 defined minimal media following a recipe used previously^1^: 1x S750 salts, 1x S750 Metals, 1% Glucose, 0.1% Glutamate + distilled H2O to final volume. 10x S750 salts: 0.5 M MOPS (adjusted to pH 7.0 with KOH), 100 mM Ammonium Sulfate, 50 mM Potassium Phosphate Monobasic (adjusted to pH 7.0). Filter sterilized. 100x S750 Metals: 0.2 M MgCl_2_, 70 mM CaCl_2_, 5 mM MnCl_2_, 0.1 mM ZnCl_2_, 100 μg/mL Thiamine HCl, 2 mM HCl, 0.5 mM FeCl_3_, dH_2_O to final volume. We performed our S750 experiments by first inoculating overnight S750 cultures from LB plates, then back diluting the following morning into fresh S750 media and allowing these new cultures to grow to exponential phase prior to starting an experiment.

### Click Labeling of Peptidoglycan Precursors

We treated exponentially growing cultures with 1mM EDA-DA (ethynyl-D-alanine D-alanine, Fisher Scientific 771410) for 20 min at 37°C. We then washed cell cultures once in PBS, before fixing cells for 20 min on ice in PBS supplemented with 3.6% paraformaldehyde. We labeled the EDA-DA alkyne group with Alexa Fluor 488 Azide (A10266) by following the Invitrogen Click-iT™ Cell Reaction Buffer Kit (Thermo Fisher, C10269), but without supplementing with BSA (to ensure optimal diffusion through the cell wall). We spotted cells onto PBS pads made with 2% agarose and imaged all samples within 18 h of click-labeling. We generated different treatment conditions by inoculating identical, prewarmed tubes of fresh growth media from a single exponentially growing culture. We generated our tunicamycin-treated samples by pre-treating for 10 min with 0.5 𝜇g/mL tunicamycin prior to EDA-DA addition, for a total of 30 min drug exposure. By contrast, we added 50 𝜇g/mL fosfomycin or 1𝜇g/mL vancomycin simultaneous with EDA-DA for a total of 20 min drug exposure.

### Protein purification and application

Protein was purified according to the following, previously published method^2^: We incubated cell cultures of bER461 in 500mL LB + 100 ug/mL Ampicillin and 35 ug/mL Chloramphenicol at 37°C with shaking until they reached an OD600 of 0.6-0.8. We then shifted these cultures to 16°C, added IPTG (Isopropyl ß-D-1-thiogalactopyranoside) to a final concentration of 0.5mM, and incubated with shaking for 18h. We centrifuged these cultures at 4°C, resuspended in Buffer A (50 mM Tris-HCl + 200 mM NaCl + 1 mM CaCl2 + 1 mM MgCl2, pH 8.0) +25mM Imidazole and lysed by disruption in an Emulsiflex-C5. After disruption, we incubated the supernatant with a column containing HisPur Ni-NTA Resin (ThermoFisher 88221), ran this column over an Imidazole gradient, and visualized samples via SDS-PAGE with stain-free labeling (BioRad 1610183). We pooled column elutions containing high protein concentrations, performed dialysis at 4°C overnight in Buffer A, concentrated our samples with Amicon Ultra Centrifugal Filters (10KDa molecular weight cutoff), and flash-froze single-use aliquots with liquid nitrogen for storage at-80°C. We calculated our protein concentration by nanodrop.

The Gram-positive cell wall imposes a substantial diffusion barrier to molecules above 15kDa^3^. At 30.1kDa, GlpQ exceeds this limit substantially, rendering it necessary to partially degrade the cell wall to allow GlpQ access to its target substrates. We accomplished this by performing all GlpQ incubations in PBS lacking divalent cations for the times listed. This incubation simultaneously arrested peptidoglycan synthesis and induced autolysis (cell wall degradation) by the cells’ native autolysins. We made the PBS used to perform these incubations according to the following recipe^4^ (10X PBS, 1L): 80g NaCl, 2g KCl, 14.4g Na_2_HPO_4_, 2.4g KH_2_PO_4_. Adjust pH to 7.4 with HCl.

### *In vivo* bocillin labeling

We grew cell cultures to early log-phase, back diluted into prewarmed LB to create multiple identical, log-phase samples, and added tunicamycin (MP Biomedical 0215002805 and Cell Signaling Technology 12819) to each sample at successive time intervals of 10 min, to a final concentration of 0.5μg/mL. Prior to lysis, we treated all cultures with bocillin-FL (Invitrogen, B13233) simultaneously to a final concentration of 1μg/mL, incubated for 2 min, then pelleted cells for cell lysate extraction. We treated cell pellets with lysis buffer (20mM Tris pH 7.5, 1mM EDTA, 10mM MgCl_2,_ 1mg/mL Lysozyme, 1mM PMSF, 10ug/mL DNAse I, 100ug/mL RNAse A). We added fresh Lysozyme, PMSF, DNAse I and RNAse A to a lysis buffer stock for each preparation. We then visualized cell lysates via SDS-PAGE (FastCast 10% Acrylamide, Bio-Rad 1610183) using a Cytiva Amersham Typhoon fluorescent imager.

### Wall teichoic acid measurement by flow cytometry

We grew cell cultures to log-phase (OD600=0.1-0.2), then took two samples from a single cell culture, washed 1X in PBS, and incubated in PBS + 40μM GlpQ or PBS + the equivalent volume of Buffer A for 15 min with rocking at 37°C. We then washed cells again 1X in PBS, and resuspended cells in PBS supplemented with 200ug/mL of Concanavalin A-AlexaFluor 647 (Invitrogen, C21421) for 10min at 37°C, with rocking. Concanavalin A is a lectin that binds glycosylation modifications along the teichoic acid polymer^5,6^. We then washed cells again 1X in PBS, then resuspended samples in PBS for flow cytometry. We performed our flow cytometry measurements with a Cytek Aurora set to 10,000 events per sample, with gains adjusted to capture the full dynamic range of our population measurements. We consistently calibrated the instrument to a low background (<10 events/second), and vigorously vortexed each sample prior to acquisition. Since we were interested in statistics of the whole cell population, we performed minimal gating (Fig. S6B). Since GlpQ preferentially cleaves non-glycosylated teichoic acids^2^, the GlpQ-mediated reduction in Concanavalin A staining for wild-type cells is likely an underestimate of GlpQ-mediated teichoic acid cleavage. We used Δ*tagE* mutants to control for non-specific wheat-germ agglutinin labeling, since TagE is responsible for teichoic acid glycosylation^7^.

### Image Analysis and Tracking

Cell tracking was performed using custom scripts written in FIJI (imagej.net), MATLAB (Mathworks, Natick, MA, USA) and Python, all of which are available online at https://github.com/felixbarber/WTA_peptidoglycan_nanostructure. We first applied the ImageJ scripts image_process_updated.ijm and alignment_updated.ijm (for timelapse imaging, e.g. Fig. 1B) or timepoint_imaging_v1.ijm (for discrete timepoint imaging, e.g. Fig. 1D) to generate the necessary tiff image file structures for downstream segmentation and tracking in MATLAB. Our MATLAB timelapse image analysis pipeline then involved the following steps: image alignment (imagealign_barber.m), background removal (to avoid spurious detections, following eraseimagepart_barber.m), followed by cell segmentation and tracking (BacTrack_barber.m). All cell segmentation was done based on phase contrast images. Once phase contrast images were analyzed to generate individual cell masks (and cell tracks in the case of timelapse imaging), we applied python scripts to analyze and plot the single cell characteristics, and to integrate fluorescence data (for our discrete timepoint data series). The applicable python scripts differ based on the purpose and plotting format for individual experiments, and a complete list of the various different pipelines for different data formats is provided in the README file within our code repository. Within these steps, we filtered our timelapse cell tracks to exclude cells tracks of 4 timepoints or shorter, to exclude non-growing (dead) cells from analysis, and to remove sections of cell tracks with a growth rate greater than two standard deviations away from the median growth rate (e.g. due to cell divisions or mis-segmentations that combined multiple objects). For our cell staining experiments, we calculated the cellular fluorescence by subtracting a background signal (obtained by normalizing a Gaussian blur of the background signal across the cell mask), then calculating the statistics listed on the pixel fluorescence values within individual cell masks. We performed spot detection (e.g. for Fig. 1D) with SciKitImage’s bloblog feature, using manually curated thresholds for each imaging condition, that were then applied consistently across all images acquired using that condition (including across different cell types).

### MreB filament tracking

We detected and tracked our MreB and Mbl filaments using custom Python scripts as follows: first, we generated our image file structures for downstream analysis using the ImageJ script tirf_processing.ijm. We then applied the python script mreb_tracking_v3_py3.py, which detected, tracked and classified filaments as follows: For filament detection, we applied SciKitImage’s bloblog feature, using manually curated thresholds to start each timelapse (that were then preserved within all timelapses acquired using the same fluorescent fusion protein and imaging condition). These thresholds were iteratively reduced throughout each step of a single time course to maintain roughly equivalent levels of spot detection. This allowed us to correct for partial photobleaching of the fluorescent signal. We tracked individual puncta across successive images (taken with a timestep of 2s) by calculating the minimum distance between all annotated puncta, subject to a maximum displacement threshold of two pixels (equivalent to a linear distance of 186nm based on our magnification and camera size, significantly greater than the average displacement of a processive individual MreB or Mbl filament of roughly 80nm). We then filtered to exclude all tracks with length 2 timepoints or less. To classify individual tracks as “ballistic”, tracks had to meet two criteria. First, we performed a linear regression on a log-log plot of distance (calculated between successive points in each single puncta track and the track’s starting position) versus time. If the slope of this linear regression (the equivalent of ½ the “diffusion exponent” within single-molecule tracking, equal here to 0.5 for diffusive behavior and 1.0 for ballistic, linear motion) was greater than 0.8, *and* if the R-value of a linear regression for a standard plot of distances versus time (a metric for the goodness of fit for linear motion) was greater than 0.9, we classified a track as ballistic.

### Cell lysis tracking

We tracked cell lysis by perfusing cells with propidium iodide at a final concentration of 2μg/mL, imaging every 5 min with 100 ms in the RFP channel at 10% power. Due to a high level of background signal following cell lysis, we annotated cell lysis events using the semi-automated custom ImageJ macros lysis_annotation_lysed_cells_updated.ijm and lysis_annotation_plain_cells_updated.ijm. These macros allowed the user to provide annotations for lysed cells and all cells respectively within each image of a propidium iodide-stained timecourse, which were then integrated and plotted using the python script lysis_timelapses.py.

### Analysis of peptidoglycan composition

Peptidoglycan was prepared from cell lysates and analysed by high-performance liquid chromatography (HPLC) as previously described^8^: cell lysates were prepared from 500 mL of cell culture by boiling the cells (100°C) for 30 min in the presence of 5% SDS to inactivate autolysins. The SDS was removed from the insoluble cell wall preparation by repeated centrifugation (130,000 x *g* - 60 min)), the crude cell wall washed with 1 M NaCl and twice with dionized water (dH_2_O), and resuspension in dH_2_O. The pellet was resuspended in 2 to 4 mL of dH_2_O, 1/3 volume of acid-washed glass beads (diameter of 0.17–0.18 mm, Sigma, München, Germany) was added, and cell walls were disrupted in a FastPrep machine (FP120, Thermo Scientific, Hemel Hempstead, UK). Broken cell walls were separated from the glass beads through filtration and collected by ultracentrifugation as above. Cell walls were incubated at 4°C with DNase A and RNase (for 2 h) and then with trypsin (18 h), and then incubated with 1% SDS at 80°C for 15 min. The cell walls were then washed with 8 M LiCl, 20 mM EDTA and water, and lyophilized. The attached wall teichoic acid was removed by treatment with 48% hydrofluoric acid (48 h at 4°C). The resulting peptidoglycan was recovered by centrifugation and washed once with 50 mM Tris/HCL pH 7.0 and then with ice-cold distilled water until a neutral pH was achieved. The peptidoglycan was digested with cellosyl (gift from Hoechst, Germany) and the resulting muropeptides recovered were reduced with sodium borohydride and separated by HPLC on a 250 × 4.6 mm 3-μm Prontosil 120-3-C_18_ AQ reversed-phase column (Bischoff, Leonberg, Germany) maintained at 52°C. A 270 min linear gradient reverse phase separation was performed using 40 mM NaPO_4_ pH 4.5, 0.003%NaN_3_ (buffer 1, called buffer A in prior work but renamed here to avoid confusion with our GlpQ protein buffer) and 40 mM NaPO_4_ pH 4.0, 20% MeOH (buffer B)^9^. Muropeptides were detected by their absorbance at 202 nm and structures were assigned by comparison to published chromatograms^9^.

### Permeability Assays

Cells were grown to log-phase and diluted 20-fold into a pre-warmed medium within the loading well of a pre-heated CellASIC B04A microfluidic flow cell. The cells incubated for an additional 30 minutes prior to loading into the imaging chamber. While imaging, fresh medium was perfused through the flow cell. The cell-trapping mechanism used by the microfluidic chips had no detrimental effect on the elongation or morphology of cell chains, as compared with cells growing on agarose pads or liquid culture^7^.

To perform membrane lysis, we exchanged the medium in the flow cell with media + 5% N-lauroylsarcosine (Sigma Aldrich L9150) for two minutes. We monitored detergent exchange using the dye AlexaFluor 647 at a final media concentration of 5 μg/mL and imaged with CY5 illumination, 640 nm excitation at 10% intensity and 50 ms exposure. After membrane lysis, the detergent in the flow cell was exchanged for media to wash the cells.

Because both tunicamycin and GlpQ are temperature sensitive, all handling steps were performed to better preserve reagent stability: tunicamycin and GlpQ were introduced into the microfluidic device concurrently with the cells. At the end of the culture back-dilution, a 5 mg/mL tunicamycin stock aliquot was thawed briefly and subjected to two successive 1:100 dilutions in LB before being loaded into the chip. GlpQ aliquots were thawed on ice for approximately 30 min and diluted in PBS immediately prior to use. Since PBS-mediated hydrolysis is sensitive to divalent cation concentration, an equal volume of Buffer A was diluted in PBS as a control.

For the experiments in Figure 3A, cells expressing mNeonGreen were imaged every minute in our GFP channel (480 nm excitation from an LED light source, Lumencor Aura Light Engine) at 100% power and 50 ms exposure. To mitigate photobleaching, no images were acquired were acquired from one minute after the onset of tunicamycin treatment to five minutes prior to detergent exchange. Wild-type and Δ*ponA* cells were exposed to 0.5 µg/mL tunicamycin for 10 minutes and 20 minutes respectively, corresponding to time points during which Rod complex activity halts in each strain.

For the experiments in Figure 3B, cells expressing mNeonGreen were imaged in our GFP channel at 100% power and 50 ms exposure: every minute for seven minutes, not acquired for eight minutes during Buffer A/GlpQ perfusion, every minute for 10 minutes, and then every five minutes for 1 hour. Cells were exposed to GlpQ or Buffer A for 15 minutes prior to membrane lysis.

For the experiment in Figure 3C, cells expressing a tetracysteine-tagged ubiquitin, we imaged in our GFP channel at 10% power and 50 ms exposure every minute for the first 10 minutes and every five minutes afterwards. To label the tetracysteine-tagged ubiquitin, we incubated cells with 1 μM FlAsH-EDT_2_ (4’,5’-bis(1,3,2-dithiarsolan-2-yl)-3’,6’-dihydroxy-spiro[isobenzofuran-1(3H),9’-[9H]xanthen]-3-one; Cayman Chemical 20704) for one hour prior to diluting the cells into the loading well, which also had 1 μM FlAsH-EDT_2_ in the medium. To perform membrane lysis and wall teichoic acid hydrolysis simultaneously, the medium in the flow cell was exchanged with PBS + 5% N-lauroylsarcosine + GlpQ (or Buffer A) for two minutes. We then perfused the cells with PBS + GlpQ (or Buffer A). As a control for non-specific binding of FlAsH, we performed the same experiment in Buffer A with uninduced cells and subtracted the mean uninduced fluorescence from the fluorescence trace of each cell.

The fluorescence from the frames during lysis were interpolated. The cellular trace was corrected for photobleaching and then normalized to the final pre-lysis frame.

To perform the photobleaching controls^3^, cells were imaged at one frame per minute during the initial seven minutes of the experiment prior to cell lysis. After membrane lysis, the cells were imaged at frame rates that were much more rapid than the rate of protein efflux.

### Limitations

our permeability assay can quantitatively assess pores in the cell wall empirically, but cannot resolve the distribution of pore sizes^3^. For example, we could not distinguish between 10 pores that allow the passage of one mNeonGreen enzyme at a time, or one pore that allows 10 mNeonGreen enzymes to pass through simultaneously. Additionally, while we controlled for the effect of the isoelectric point (pI) of the probe, we have not yet explored whether there are other electrical properties of the probe that influence permeability through the wall (for example, the net charge or distribution of acidic/basic residues throughout the protein).

### Permeability photobleaching correction

Population-averaged fluorescence traces were fit to a single exponential:

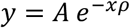

where y = normalized fluorescence, A is the initial post-lysis normalized fluorescence value, 𝜌 = the photobleaching constant, and x is the frame number (Figure S7, S9).

For mNeonGreen, the mean background value is subtracted from the mean fluorescence of each cell and normalized by the fluorescence of the final pre-lysis frame.

For FlAsH, the mean background value and mean autofluorescence is subtracted from the mean fluorescence of each cell. The mean autofluorescence was obtained by measuring the fluorescence of uninduced cells incubated with FlAsH-EDT_2_.

To correct for photobleaching, the expected loss of fluorescence due to photobleaching was calculated,

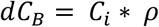

where C_B_ = bleached fluorophore, C_i_ = measured fluorescence in the *i*th frame, and r = photobleaching constant (obtained from the photobleaching control). The total loss of fluorescence (*d*C_T_) in each frame is adjusted to account for the expected loss of fluorescence due to photobleaching,

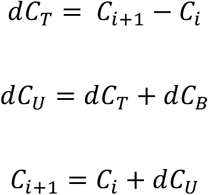

where dC_U_ is the change in unbleached fluorophore.

### Transmission Electron Microscopy

We performed transmission electron microscopy on thin sections of formaldehyde-fixed freeze-substituted *B. subtilis* cells as follows: we grew cultures of wild-type and Δ*ponA* cells to log-phase, then back diluted into either fresh LB medium or LB supplemented with 0.5𝜇g/mL tunicamycin for 30 min. We then pelleted cells by centrifugation for 4 min at 3500 RPM, washed 1X in PBS, and resuspended in 1 mL freshly made LRR fixative solution^10^ (0.5 mL 0.15% ruthenium red, 125 μL 16% formaldehyde, 0.0155 g lysine acetate (Sigma); distilled water to 1 mL).

Samples were then subjected to high pressure freezing and freeze substitution. 3 mm planchettes 100 mm deep were lightly coated with hexadecene before being filled with approximately 1.2mL of fixed sample. An absorbent filter paper was used to soak out extra liquid, and then the sample was allowed to dry slightly for ∼ 5 min. The hats were then sealed in the planchette holder for high-pressure freezing (Leica ICE High Pressure Freezing Platform, Leica Microsystems). The frozen samples were immediately transferred into liquid nitrogen and then into cryovials containing freeze substitution solutions (2 mL cryovials (Nalgene) containing 2% (wt/vol) osmium tetroxide (OsO_4_) and 0.1% (wt/vol) uranyl acetate in anhydrous acetone with 0.075% (wt/vol) ruthenium red and 2% H_2_O) at liquid nitrogen temperature. The samples were brought into a Leica AFS2 EM freeze substitution unit (Leica Microsystems) and left in the - 90°C for 79 hours. Since the acetone:osmium mixture liquifies at-90°C, the hats were slowly submerged in the freeze substitution media. The temperature of the unit was raised 5°C per hour to –60°C and incubated for 12 hours, then to –30°C for an additional 6 hours, and finally to a temperature of 0°C for 6 hrs.

After removal from freeze substitution unit, samples were washed in pure ethanol 3 x 1 hour on ice to rinse out osmium. They were then washed with 1:1 100% ethanol:LR White resin for 2 hours on ice, 1:2 100% ethanol:LR White resin overnight at 4°C, pure LR White resin for 8 hours on ice, and finally pure LR White resin overnight at 4°C. Samples were then embedded in gelatin capsules filled with LR White resin at room temperature and polymerized for 48 hours at 55-60 °C. Thin sections were cut onto 200 mesh grids and counterstained with 4% aqueous uranyl acetate for 5 min. Stained grids were examined using a JEOL1400 Flash transmission electron microscope (Japan) and photographed with a Gatan Rio 16 camera (Gatan Inc. Pleasanton, CA). All chemicals and EM grids are purchased from Electron Microscopy Sciences, Hatfield, PA.

Note: TEM sample preparations of wild-type and Δ*ponA* mutants were performed as separate batches, with Δ*ponA* mutant samples being exposed to LRR fixative solution for up to 10 min longer than wild-type cells immediately following lysis. We attribute this difference to the darker cell wall appearance of Δ*ponA* mutants relative to wild-type cells, making it difficult to compare absolute cell wall thickness between these cell types.

## Data Availability

All datasets presented herein will be made publicly available.

## Code Availability

All code used herein is publicly available at: https://github.com/felixbarber/WTA_peptidoglycan_nanostructure

## Acknowledgements

ERR and FB were supported by NSF-CAREER grant 2047404. WV received funding from the UK Biotechnology and Biological Sciences Research Council (BBSRC) (BB/W013630/1). ZA was supported by NIH grant R35GM143057. We thank Ethan Garner and the Garner Lab (both past and present) for generously sharing strains; we have particularly benefitted from strains built by prior Garner Lab members Mrinal Kapoor, Mike Dion, Yingjie Sun and Sean Wilson. We also thank the Brown lab for sharing their GlpQ expression construct. We thank Dr. Daniela Vollmer for preparation of peptidoglycan. We thank NYULH Microscopy Lab Alice Liang, Jason Yin and Jason Liang (partially supported by NYU Cancer Center Support Grant NIH/NCI P30CA016087) for consultation and timely preparation of the electron microscopy works.

## Author Contributions

FB, ZA, ZY, JB and LPL performed the experiments. FB, ZA, ZY, JB and LPL analyzed the data. FB and ER wrote the manuscript.

## Extended Data: Tables

**Table S1:**
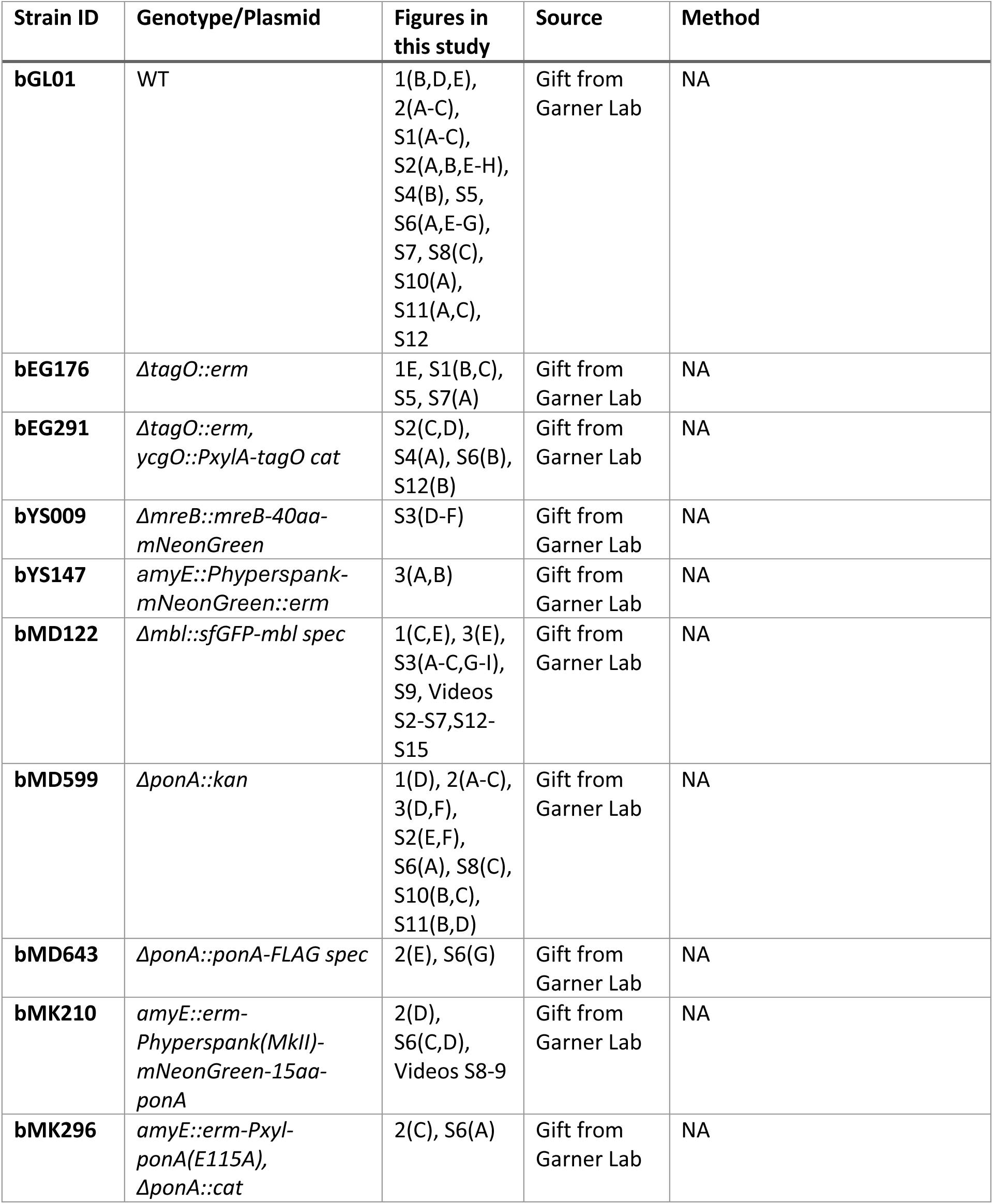
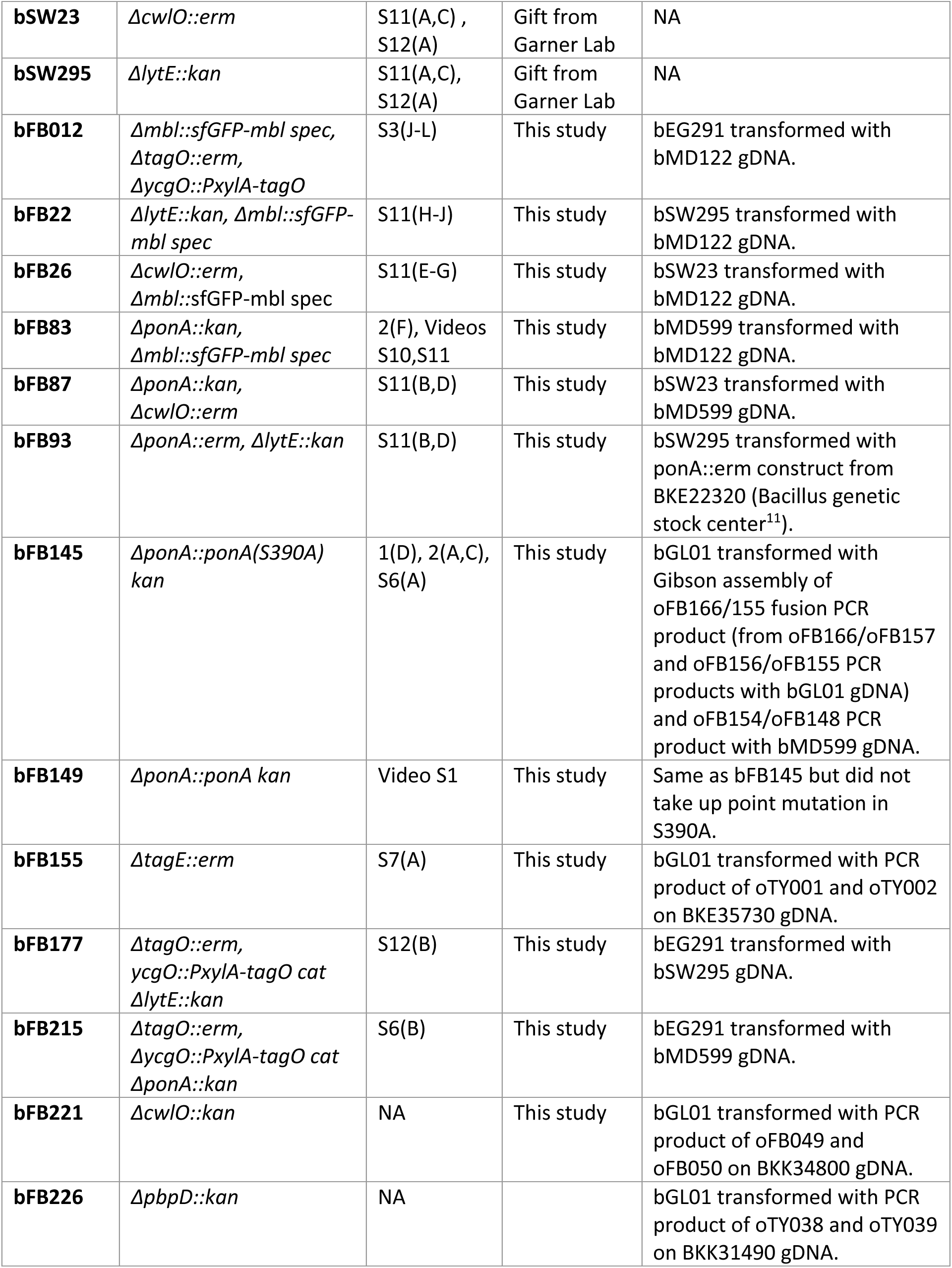
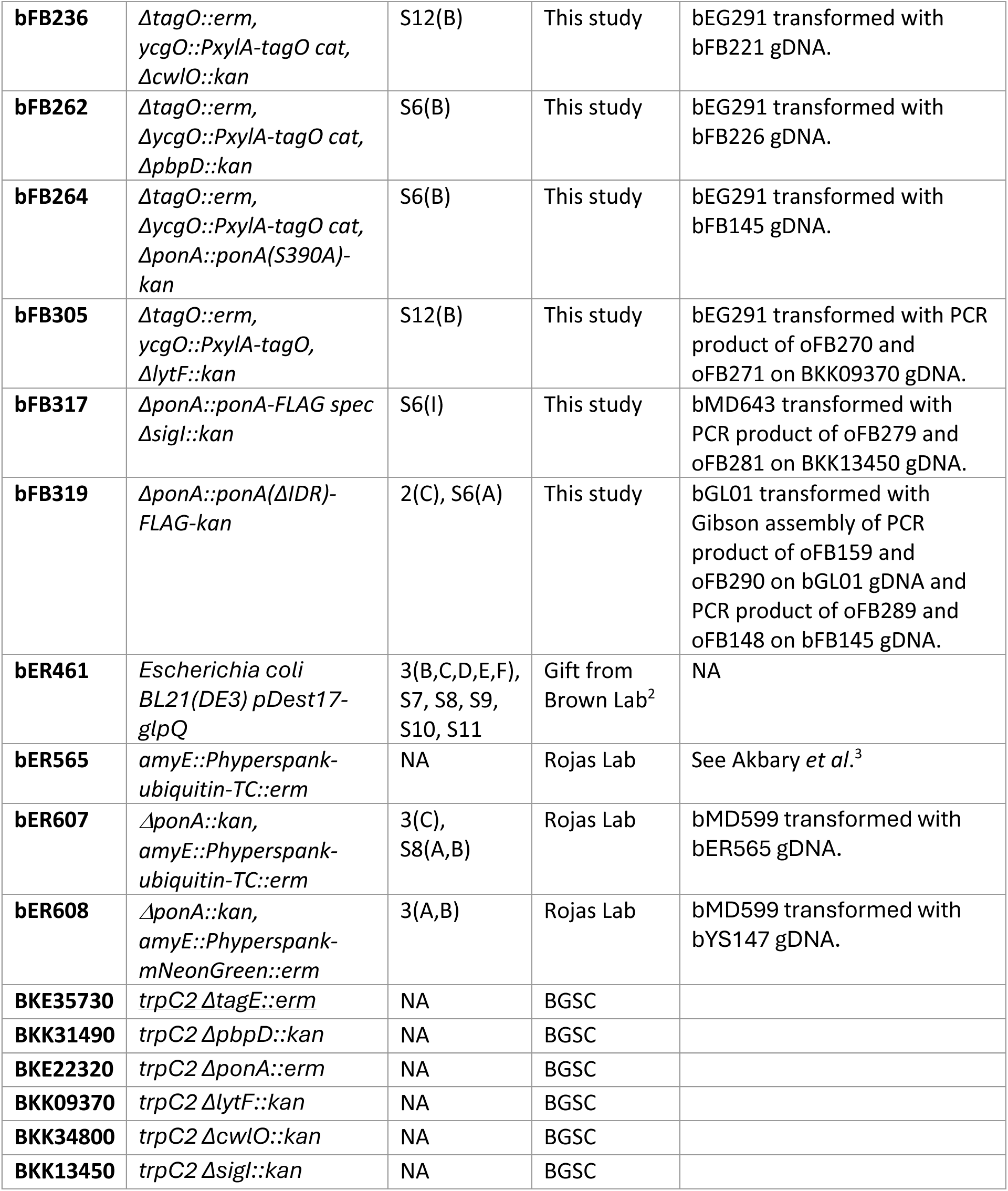
List of strains used in this study.

**Table S2:**
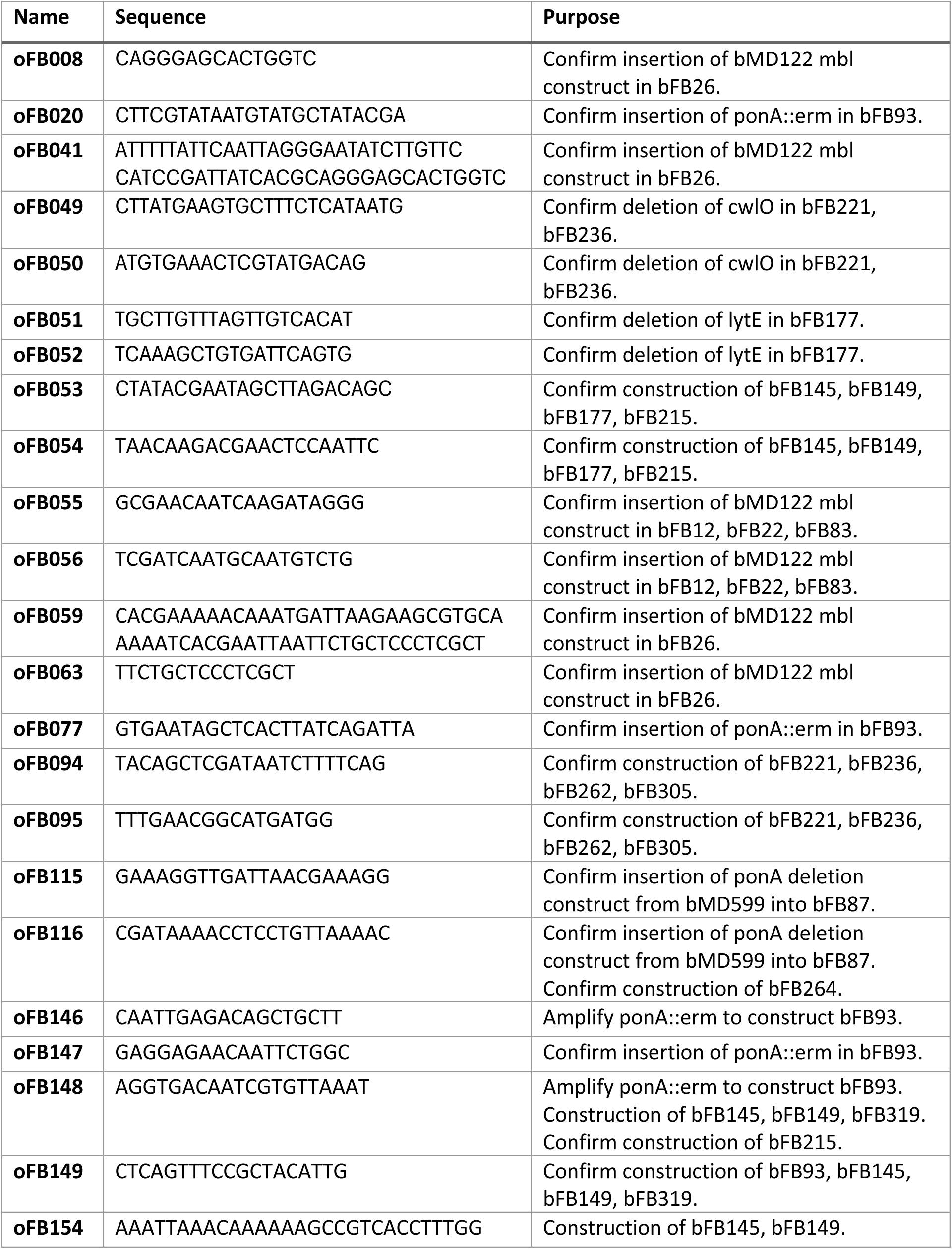
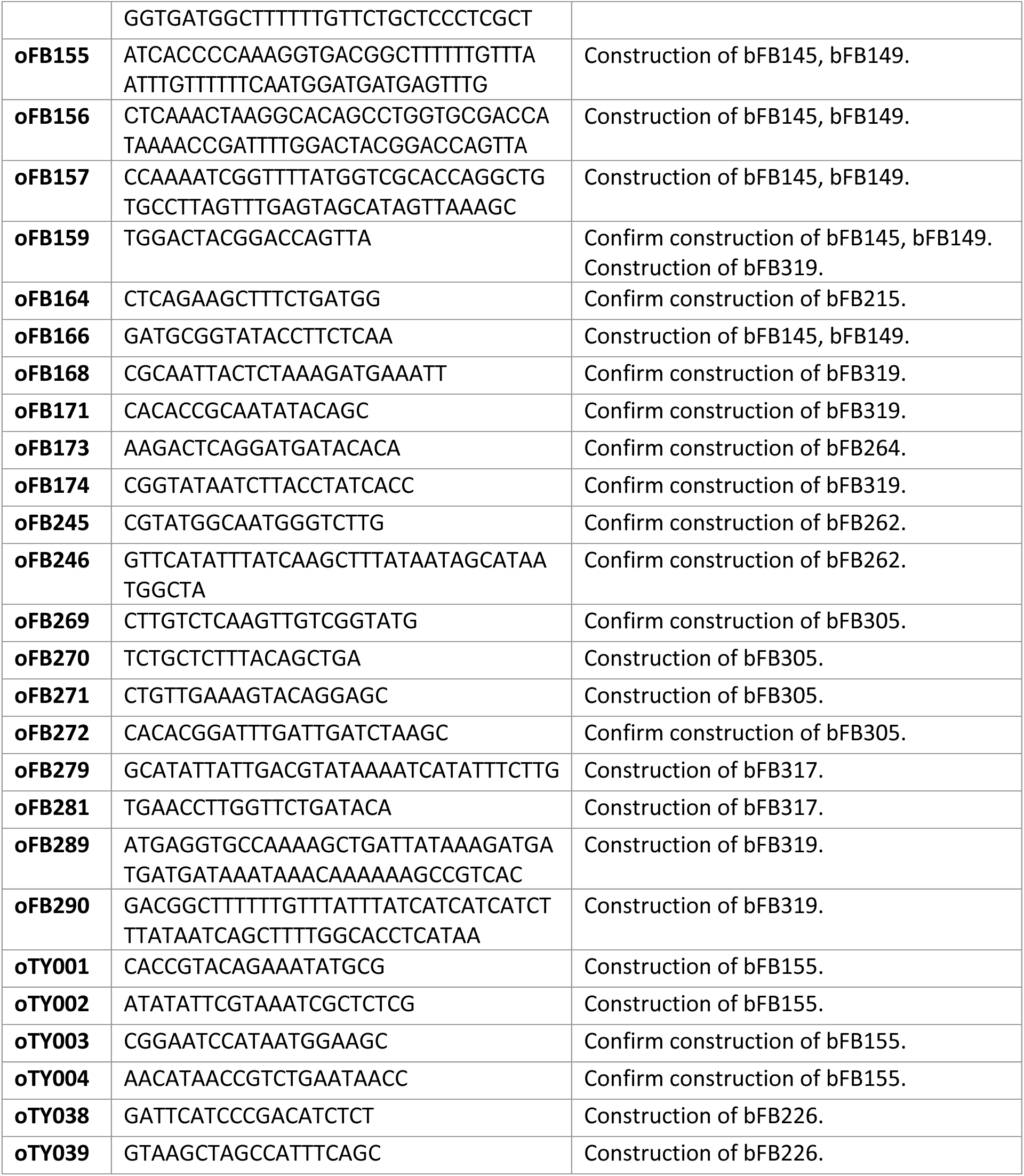
List of primers used in this study.

## Extended Data: Figures

**Figure S1:**
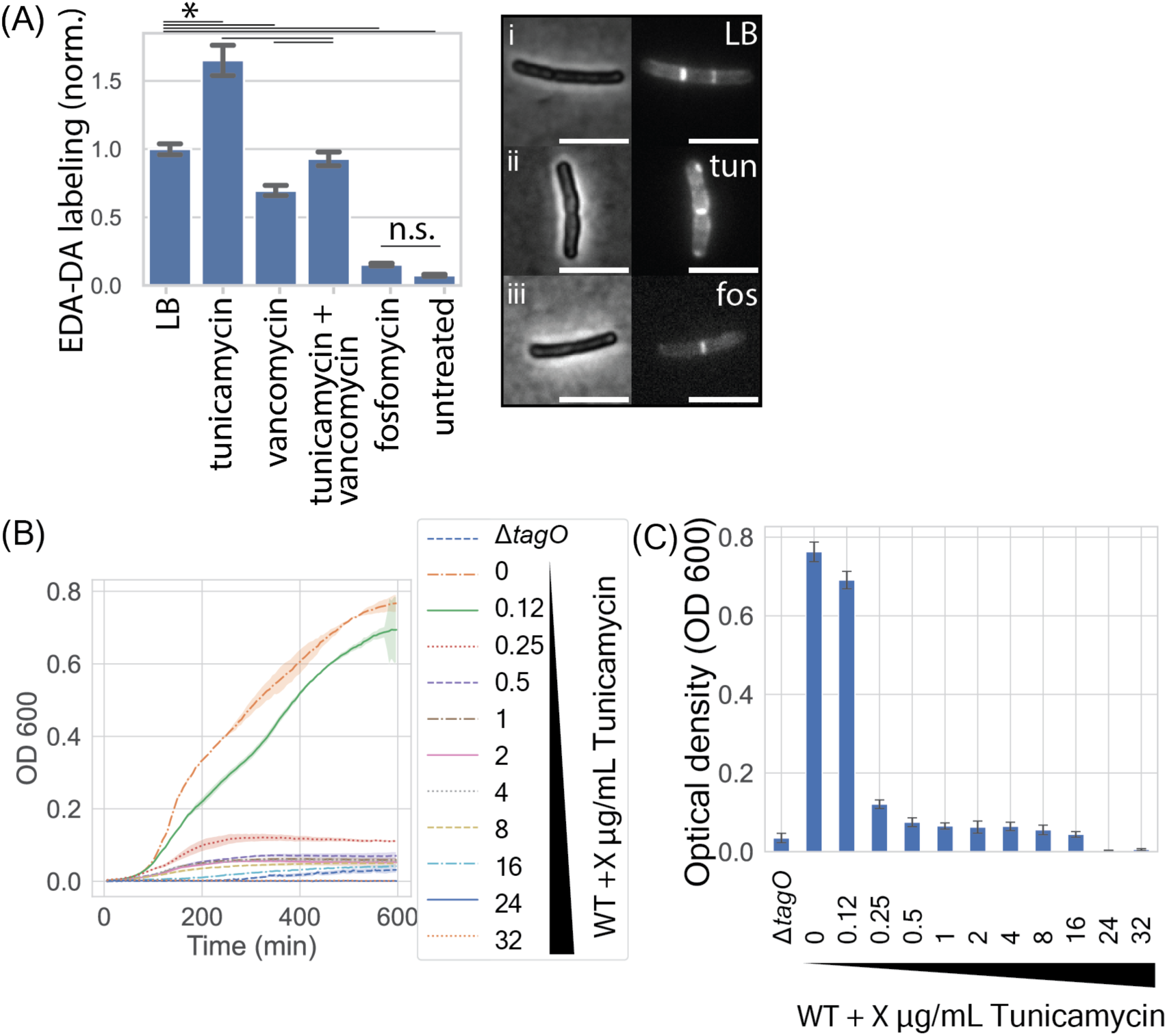
(A) Low doses of tunicamycin do not decrease peptidoglycan precursor synthesis. We pulsed exponentially growing cells with the dipeptide D-amino acid EDA-DA (ethynyl-D-alanine D-alanine), which is incorporated into peptidoglycan precursors cytosolically^12^, before fixing and “clicking” with AlexaFluor 488 azide (*Methods*). Plot shows average intensity of AF488 labeling for cells co-treated with the drugs shown, normalized against staining for cells grown in LB supplemented with EDA-DA alone. Staining increased during tunicamycin treatment (relative to EDA-DA alone), and during co-treatment with both tunicamycin and vancomycin (relative to co-treatment with vancomycin), consistent with a rerouting of precursors to peptidoglycan synthesis during TagO inhibition. Staining decreased only partially for vancomycin treatment relative to LB alone, consistent with AF488 labeling both nascent peptidoglycan and lipid II. Fosfomycin treatment arrests peptidoglycan precursor synthesis at the inner face of the plasma membrane so is a negative control. Untreated cells were not exposed to EDA-DA but were clicked with AF488 azide to control for off-target AF488 labeling. Analysis was performed over 1,533 cells (LB, 5 experimental replicates), 1,063 cells (tunicamycin, 5 experimental replicates), 810 cells (tunicamycin + vancomycin, 3 experimental replicates), 1,081 cells (vancomycin, 4 experimental replicates), 555 cells (fosfomycin control, 4 experimental replicates), and 473 cells (untreated, 4 experimental replicates). All differences between conditions were significant (one-way ANOVA followed by Tukey’s honestly significant difference (HSD) post-hoc test, P<0.01), with the exception of comparisons between fosfomycin-untreated and between LB-tunicamycin+vancomycin. Inset: phase contrast and fluorescence micrographs of cells from (i) LB, (ii) tunicamycin and (iii) fosfomycin conditions. Fluorescence micrographs are identically saturated. Scale bars 5µm. **(B-C) Low tunicamycin concentrations inhibit growth to precisely the same extent as deletion of *tagO.* (B)** Growth curves of cells grown with increasing tunicamycin concentrations in bulk culture. **(C)** Saturating culture density plotted for data from (A). Representative data from one of three biological replicates, with 4 technical replicates per condition. Error bars show 95% confidence intervals based on bootstrap analysis. Complete growth arrest consistently occurs at either 16µg/mL or 24µg/mL tunicamycin.

**Figure S2:**
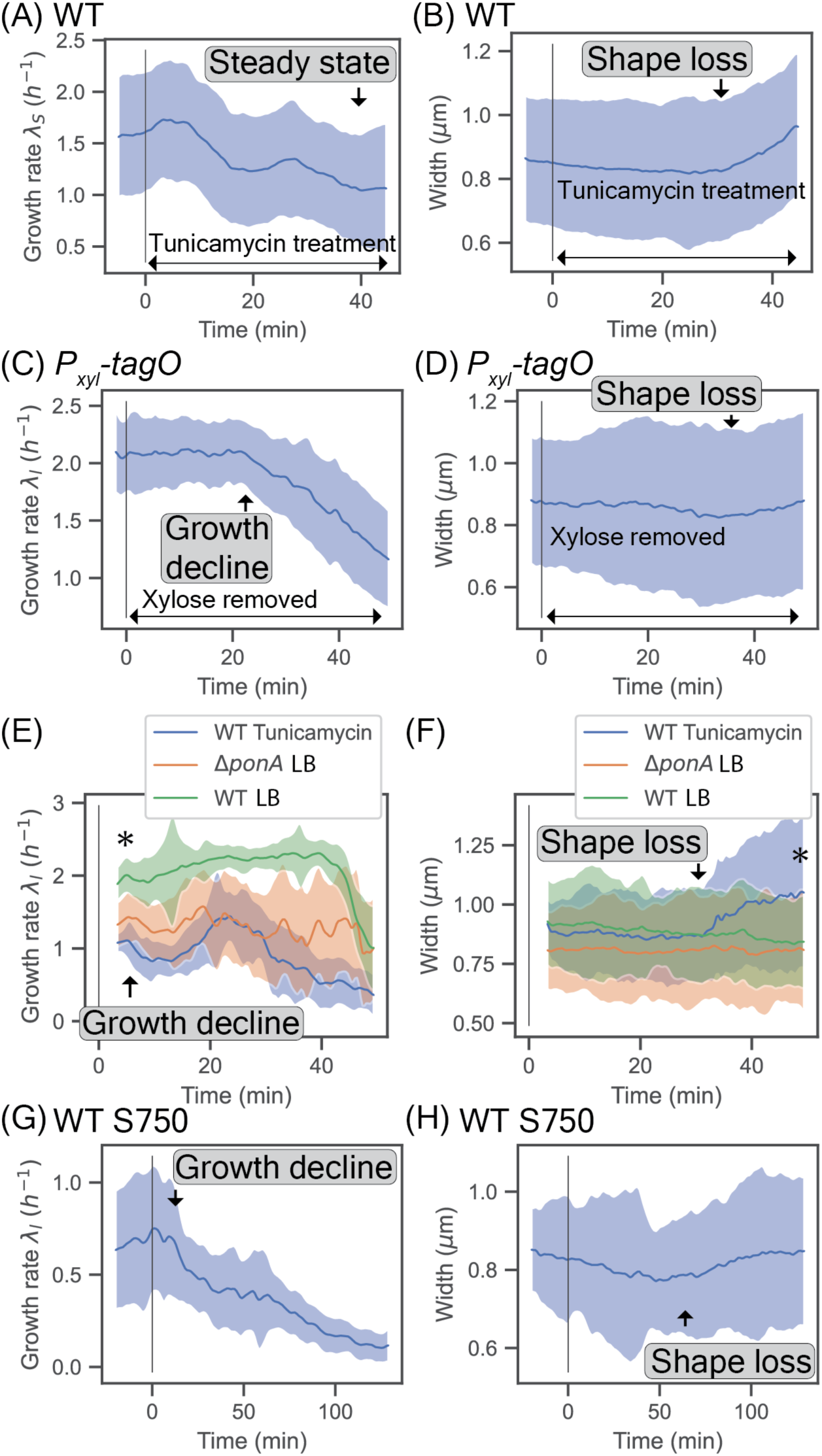
(A) Cell surface area growth rate 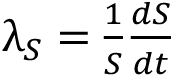 for wild-type cells during 0.5𝜇g/mL tunicamycin treatment. Analysis performed over 1,927 discrete cell tracks from 3 biological replicates. In all plots, solid lines show smoothed population medians, error bars show standard deviations across cells. Black lines show onset of perturbation (tunicamycin treatment, inducer depletion or cell spotting onto agarose pads). **(B)** Cell width of wild-type cells during 0.5µg/mL tunicamycin timecourse. Solid line shows smoothed population median, error bars show standard deviation. Black lines show onset of tunicamycin treatment. Increase in width corresponds to gradual loss of cell shape. Analysis performed over 3,737 discrete cell tracks from 3 biological replicates. **(C-D)** Cell properties during growth response of *P_xylA_-tagO* cells to xylose depletion. **(C)** Cell length growth rate. Analysis performed over 1,217 discrete cell tracks from 3 biological replicates. **(D)** Cell width. Analysis performed over 1,846 discrete cell tracks from 3 biological replicates. **(E-F)** Cell growth and shape response to tunicamycin treatment on agarose pads. **(E)** Cell length growth rate. Analysis performed over 204 discrete cell tracks from 4 biological replicates (WT LB), 160 discrete cell tracks from 3 biological replicates (WT tunicamycin) and 93 discrete cell tracks from 3 biological replicates (Δ*ponA* LB). Growth rate during tunicamycin-treatment shows immediately slower growth, with oscillations, followed by a sustained secondary decline at approximately 30 min (coincident with shape loss, see (F)). WT cells grown in LB alone showed an abrupt growth decline at 45 min with no shape loss. This late-stage decline was not observed for the slower-growing Δ*ponA* cells, consistent with local nutrient depletion in these roughly 1mm-thin agarose pads. WT LB and WT Tunicamycin time courses show statistical significance based on Student’s T-test, P<0.01, calculated at first measured timepoint. **(F)** Cell width. Analysis performed over 535 discrete cell tracks from 4 biological replicates (WT LB), 432 discrete cell tracks from 3 biological replicates (WT tunicamycin) and 214 discrete cell tracks from 3 biological replicates (Δ*ponA* LB). WT LB and WT Tunicamycin time courses show statistical significance based on Student’s T-test, P<0.01, calculated at last measured timepoint (63 min). **(G-H)** Time course of 0.5µg/mL tunicamycin exposure for WT cells grown in S750 media. **(G)** Cell length growth rate. Analysis performed over 249 discrete cell tracks from 2 biological replicates. **(H)** Cell width. Analysis performed over 307 discrete cell tracks from 2 biological replicates.

**Figure S3:**
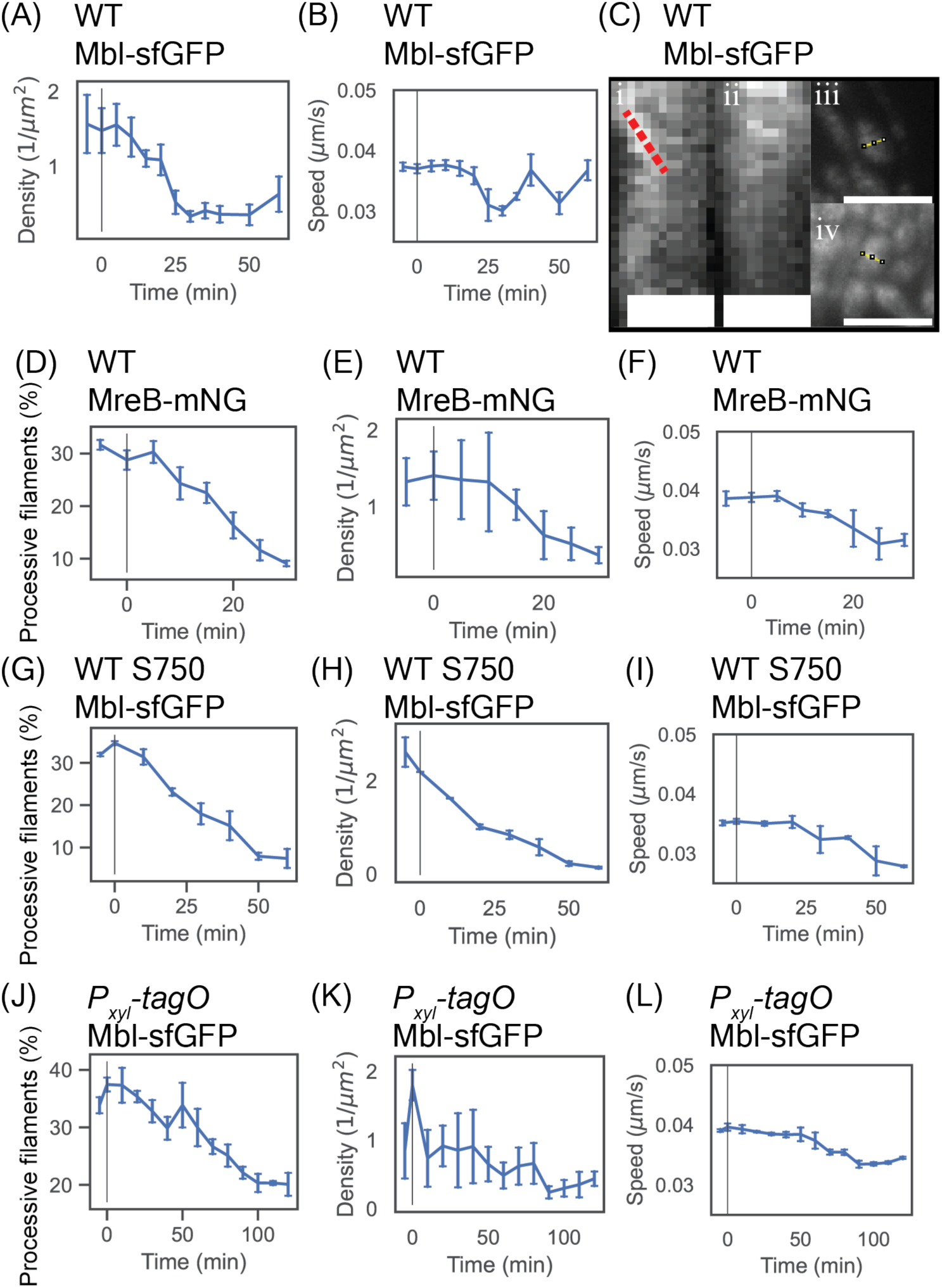
(A-B) Mbl-sfGFP activity time course for wild-type cells during 0.5µg/mL tunicamycin treatment. Analysis performed on 62,901 discrete filament tracks from 4 biological replicates. All error bars show 95% confidence intervals across biological replicates based on bootstrap analysis. **(A)** Spatial density of processive filaments. **(B)** Average speed of processive filaments. **(C)** (i-ii) Representative kymographs from TIRF imaging of Mbl-sfGFP cells after 60 min tunicamycin treatment. Red dotted line follows a processive Mbl filament. Scale bars 1𝜇m. (iii-iv) TIRF micrographs showing whole cells during a single timepoint of the kymographs in (i-ii) respectively. Yellow lines show kymograph traces. Scale bars 5µm. **(D-F)** MreB-mNeonGreen activity time course for wild-type cells during 0.5µg/mL tunicamycin treatment. Analysis performed on 30,805 discrete filament tracks from 3 biological replicates. **(D)** Percentage of processive filaments. **(E)** Spatial density of processive filaments. **(F)** Average speed of processive filaments. **(G-I)** Mbl-sfGFP activity time course for wild-type cells grown in S750 media during 0.5µg/mL tunicamycin treatment. Analysis performed on 9,859 discrete filament tracks from 2 biological replicates. **(G)** Percentage of processive filaments. **(H)** Spatial density of processive filaments. **(I)** Average speed of processive filaments. **(J-L)** Mbl-sfGFP activity time course of of *P_xylA_-tagO* cells during xylose depletion. Analysis performed on 55,542 discrete filament tracks from 3 biological replicates. **(J)** Percentage of processive filaments. **(K)** Spatial density of processive filaments. **(L)** Average speed of processive filaments.

**Figure S4.**
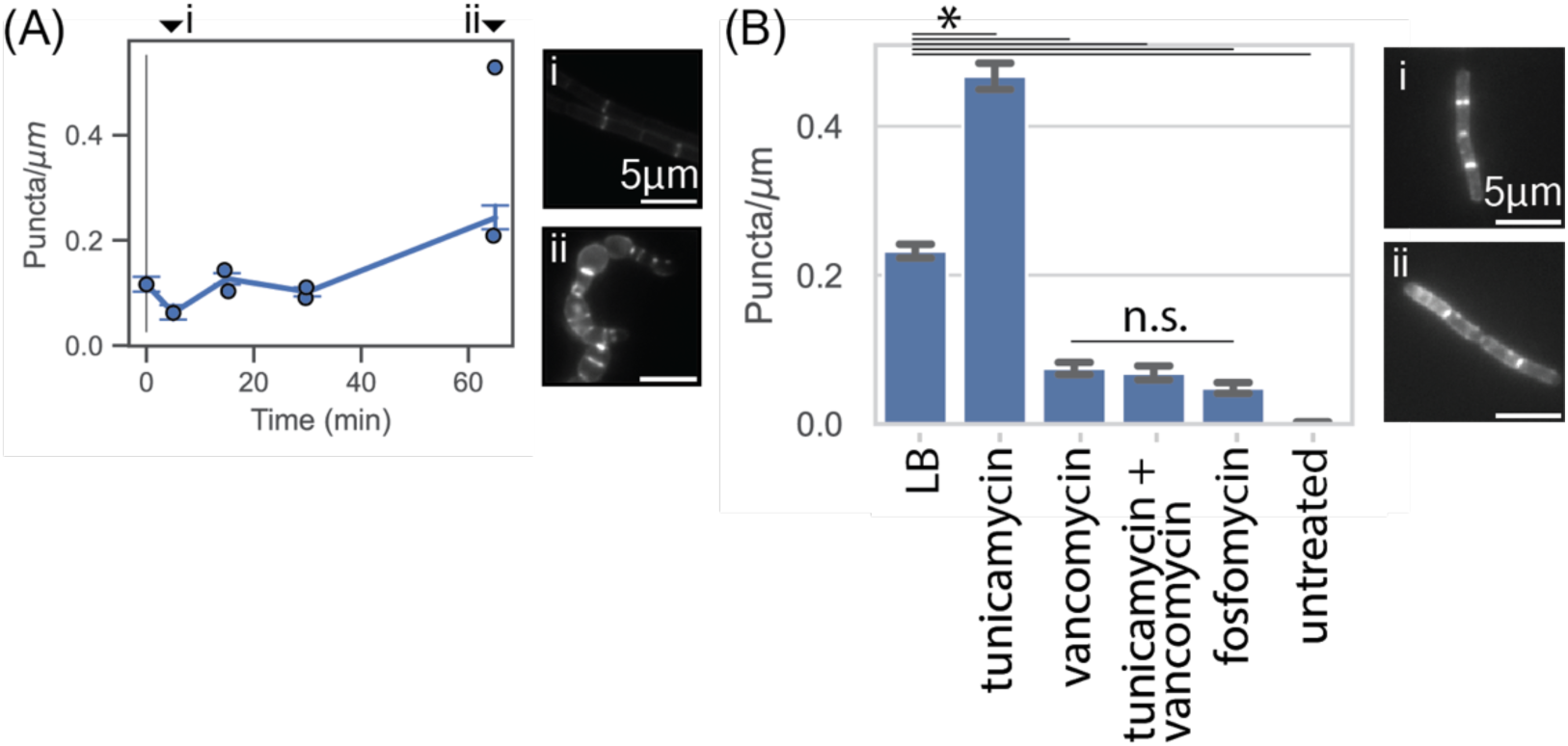
(A) Mean fluorescent puncta per cell (normalized to cell length) for *P_xylA_-TagO* cells during xylose depletion, labeled with fluorescent D-amino acids. Black line shows xylose washout. Error bars show 95% confidence intervals. Analysis performed over 680 segmentations from 2 biological replicates. Inset shows micrographs at (i) 5 min and (ii) 65 min following xylose depletion. Micrographs are identically saturated across timepoints. **(B)** Mean EDA-DA AF488 click-labeled puncta per cell (normalized to cell length) for wild-type cells during co-treatments with antibiotics shown. Analysis was performed over 1,533 cells (LB, 5 experimental replicates), 1,063 cells (tunicamycin, 5 experimental replicates), 810 cells (tunicamycin + vancomycin, 3 experimental replicates), 1,081 cells (vancomycin, 4 experimental replicates), 555 cells (fosfomycin control, 4 experimental replicates), and 473 cells (unstained, 4 experimental replicates). All differences between conditions were significant (one-way ANOVA followed by Tukey’s HSD post-hoc test, P<0.01) with the exception of comparisons between vancomycin, tunicamycin+vancomycin and fosfomycin. Inset: fluorescent micrographs of AF488 labeling in cells cultured with (i) LB alone and (ii) tunicamycin. Micrographs are identically saturated across conditions. Scale bars 5𝜇m.

**Figure S5:**
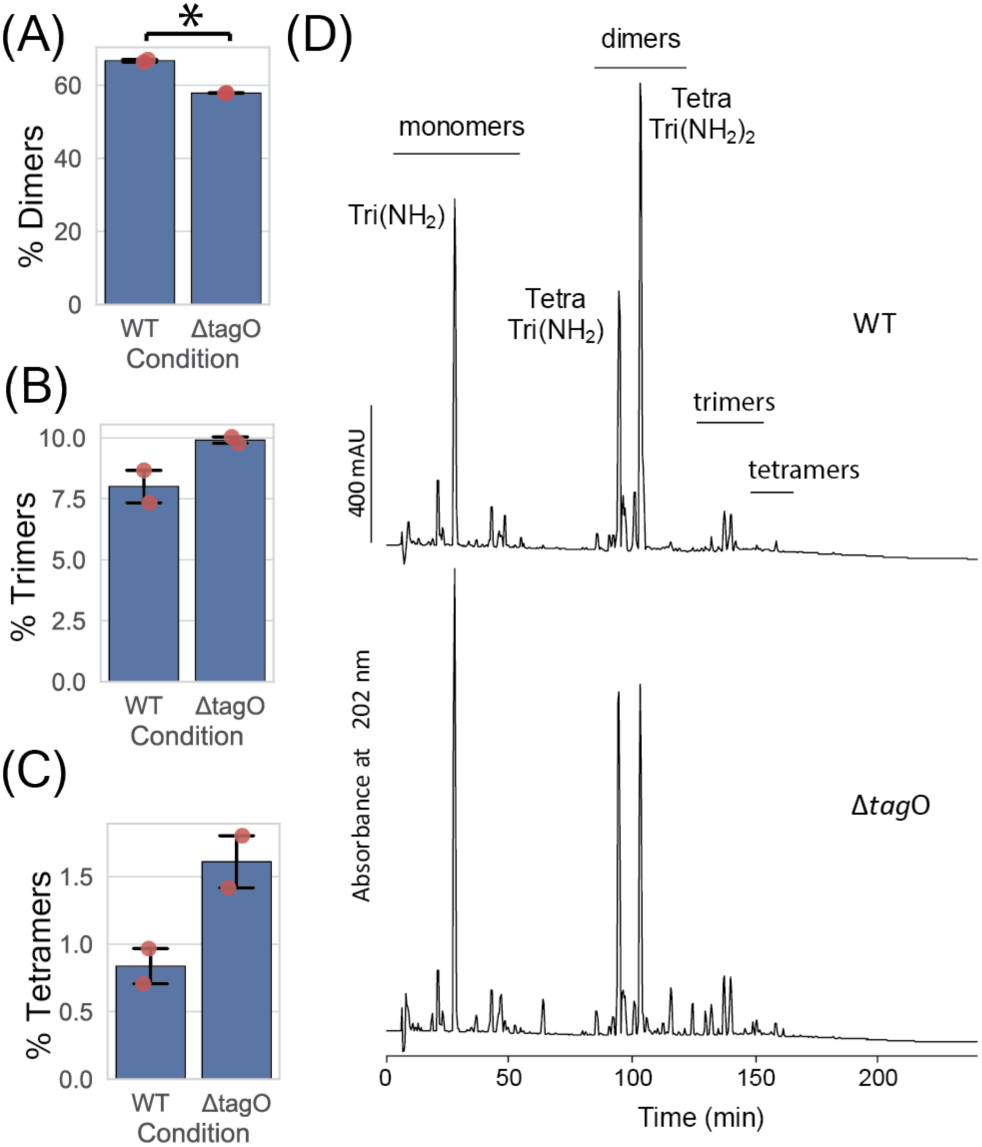
(A-C) Percentage peptidoglycan subunits in **(A)** dimers, **(B)** trimers and **(C)** tetramers. Two biological replicates per strain. Mean values calculated across biological replicates. Error bars show 95% confidence intervals. Individual datapoints shown in red. Statistical significance calculated using Student’s T-test, P<0.05. Differences for (B) and (C) are not statistically significant by this metric but reflect visible shifts in high time peaks that are consistent between samples (see (D)). **(D)** High-performance liquid chromatograms of reduced muropeptide samples isolated from wild-type and Δ*tagO* cells. Elution regions of the uncross-linked monomers and cross-linked dimers, trimers and tetramers are shown, and the three main muropeptides are labelled. Tri(NH_2_), GlcNAc-MurNAc(red)-L-Ala-D-iGlu-mDap (1 amidation); TetraTri(NH2), GlcNAc-MurNAc(red)-L-Ala-D-iGlu-mDap-D-Ala-mDap-D-iGlu-L-Ala-(GlcNAc)MurNAc(red) (1 amidation); TetraTri(NH2)2, GlcNAc-MurNAc(red)-L-Ala-D-iGlu-mDap-D-Ala-mDap-D-iGlu-L-Ala-(GlcNAc)MurNAc(red) (2 amidations). GlcNAc, N-acetylglucosamine; MurNAc(red), N-acetylmuramitol; L-Ala, L-alanine; D-iGlu, D-isoglutamic acid; mDap, meso-diaminopimelic acid; D-Ala, D-alanine.

**Figure S6:**
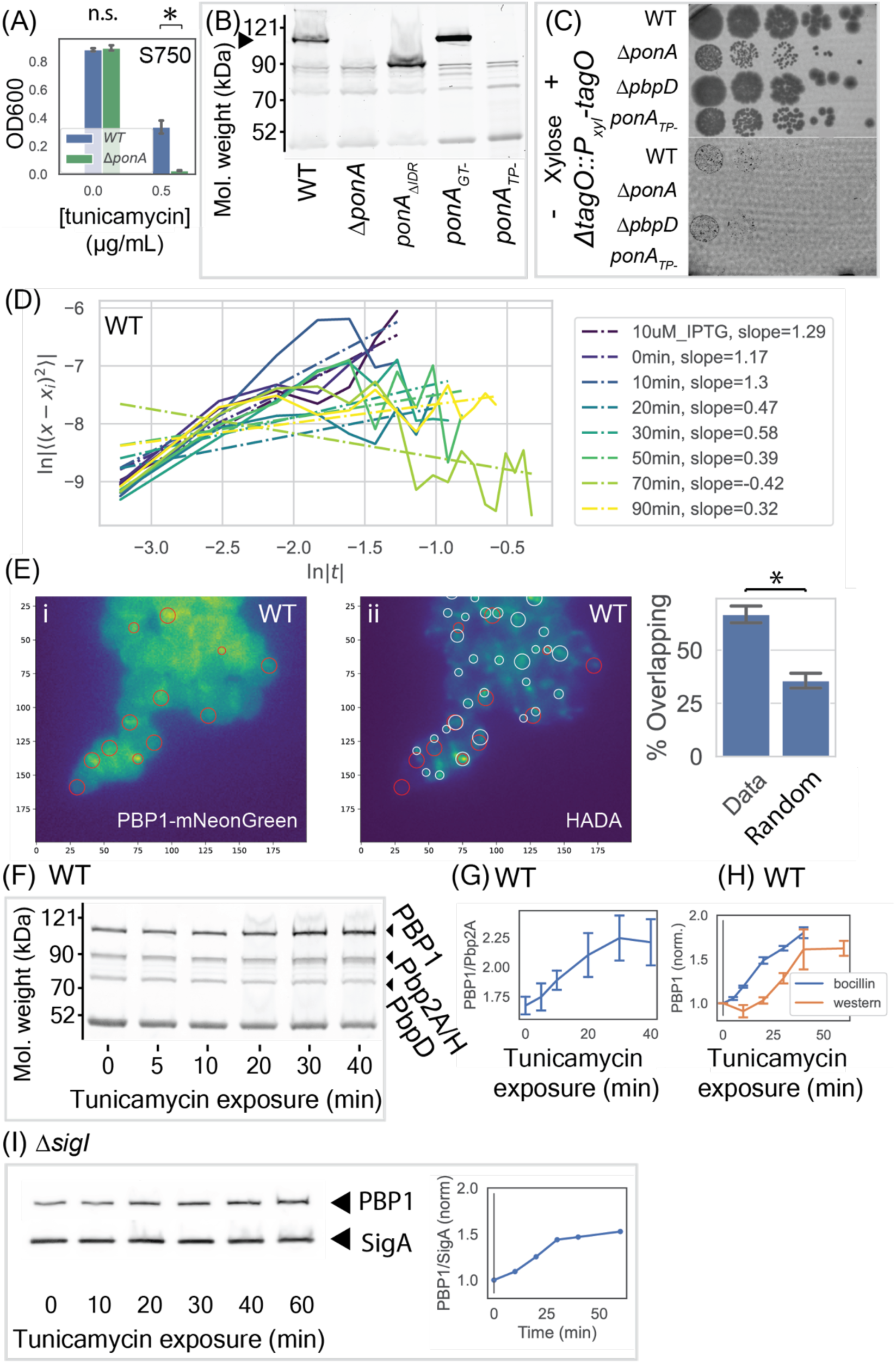
(A) Maximum OD for wild-type and Δ*ponA* cells with and without a low dose of tunicamycin, grown in bulk culture in S750 minimal media. Data shown from one representative biological replicate out of three, with four technical replicates per biological replicate. Error bars show 95% confidence intervals based on bootstrap analysis. Statistical significance calculated using Student’s T-test, P<0.01. **(B)** SDS-PAGE on cell lysates from cultures incubated with fluorescently conjugated bocillin. Black arrow shows wild-type PBP1, which is present in wild-type and *ponA_GT-_* strains, shifted in *pon*𝐴_-_*_IDR_* cells, absent in Δ*ponA* cells, and not labeled in *ponA_TP-_* cell lysates. **(C)** Spot assays for WT, Δ*ponA*, *ponA_TP-_* and Δ*pbpD* mutant strains in a *P_xylA_-tagO* background, grown in the presence and absence of 30mM xylose. **(D)** log-log plot of mean-squared displacement vs. time for single molecule tracking of PBP1 puncta, throughout a time course of tunicamycin treatment. Slopes correspond to the diffusion exponent 𝛼 in Fig. 2D of the main text. Data plotted from one representative biological replicate of three. **(E)** Co-localization of fluorescent puncta within epifluorescent imaging of (i) PBP1-mNeonGreen expressed at a low level alongside the native copy of *ponA*, and (ii) fluorescent labeling with HADA. Cells were imaged following 90 minutes exposure to tunicamycin and a 5-minute incubation with HADA. White circles show computationally detected HADA puncta, red circles show the same for PBP1. Inset: the percentage of PBP1 puncta that overlap with HADA puncta, both for our measured data and for the same PBP1 puncta when distributed randomly throughout the cell masks. Analysis performed on 3,320 PBP1 puncta and 6,253 HADA puncta across two biological replicates. Error bars show 95% confidence intervals based on bootstrap analysis, statistical significance calculated by Student’s T-test, P<0.01. **(F)** SDS-PAGE of *B. subtilis* whole cell lysates shows bocillin-PBP binding during tunicamycin treatment. Arrows show PBP1, PBP2A/PBP2B and PbpD bands. Cells were treated with fluorescent bocillin for two minutes prior to cell lysis. One representative sample shown from 3 biological replicates. **(G-H)** Quantification of (F), showing the ratio of **(G)** PBP1 to PBP2A/PBP2B and **(H)** PBP1 to PbpD intensities (normalized to value at 0 min; PbpD staining showed no sustained trend during tunicamycin treatment). Error bars show standard error of the mean across 3 biological replicates. Western blot data reproduced from Fig. 2E. **(I)** Western blot analysis of PBP1 levels during 0.5µg/mL tunicamycin treatment of a Δ*sigI* mutant. Left: Western blot bands for PBP1-FLAG and identically loaded SigA control. Loading volumes normalized based on OD600 readings. Right: Quantification of PBP1 staining, normalized by SigA stain and measured relative to values at 0min tunicamycin treatment. One biological replicate.

**Figure S7:**
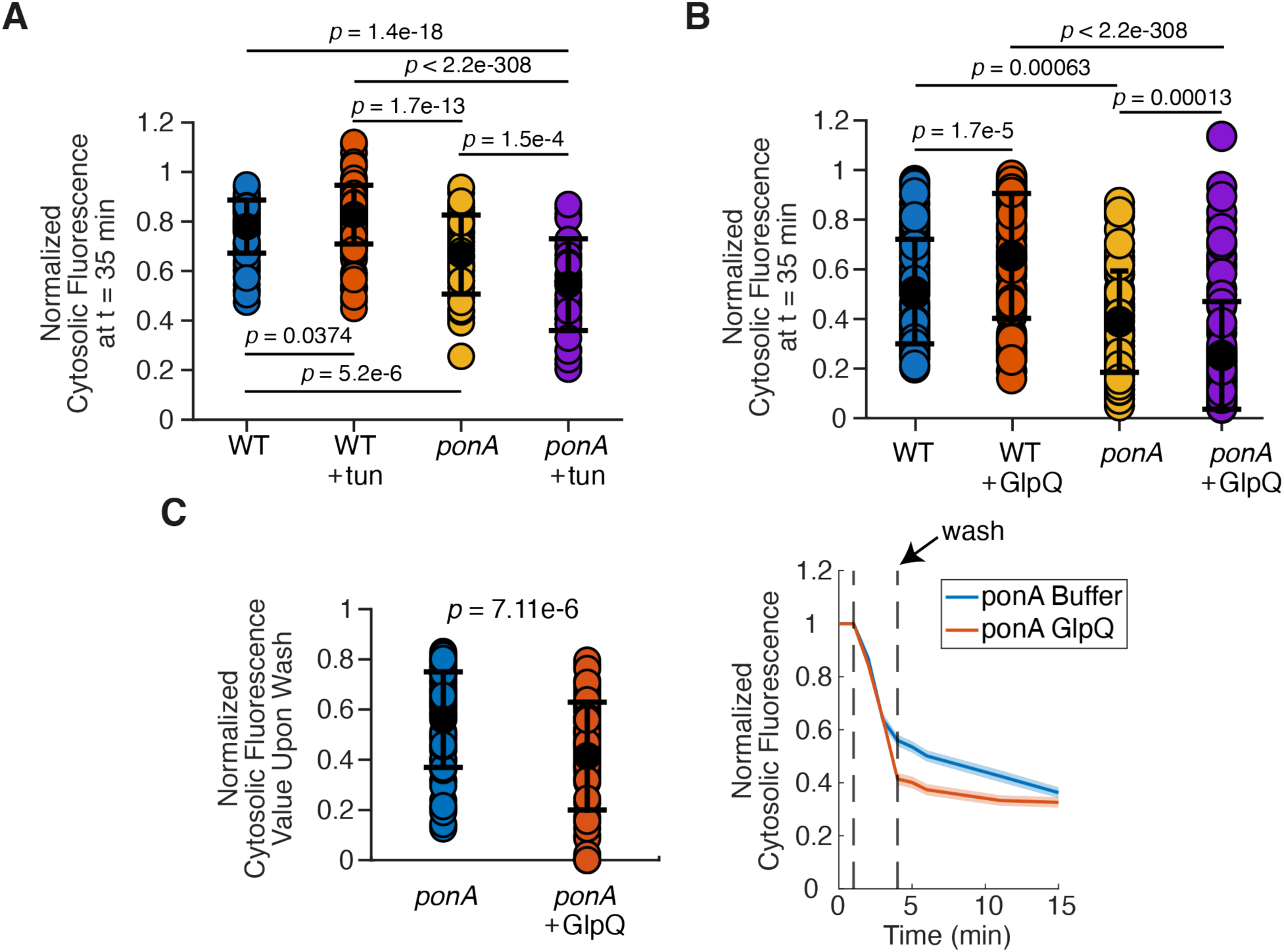
Statistical significance and decay statistics for cell wall permeability assay. (A) mNeonGreen fluorescence at t=35 min for wild-type and Δ*ponA* cells either treated with tunicamycin or untreated. Statistical significance determined via one-way ANOVA (*p* = 1.9e-27) followed by Tukey’s HSD post-hoc test for multiple comparisons (p-values shown in figure). (B) mNeonGreen fluorescence at t=35 min for wild-type and Δ*ponA* cells either treated with GlpQ in PBS or PBS alone. Statistical significance determined via one-way ANOVA (*p* = 8.7e-32) followed by Tukey’s HSD post-hoc test for multiple comparisons. All p-values are shown in the figure, except for wild-type untreated vs. *ponA* + GlpQ (*p* = 2.5e-18) and WT + GlpQ vs *ponA* untreated (*p* = 9.9e-15). (C) mUbiquitin fluorescence upon wash (as indicated by the arrow on the fluorescence vs time graph, right) following incubation of Δ*ponA* cells with detergent +/- GlpQ. Statistical significance calculated using Student’s T-test with *p* = 7.11e-6.

**Figure S8:**
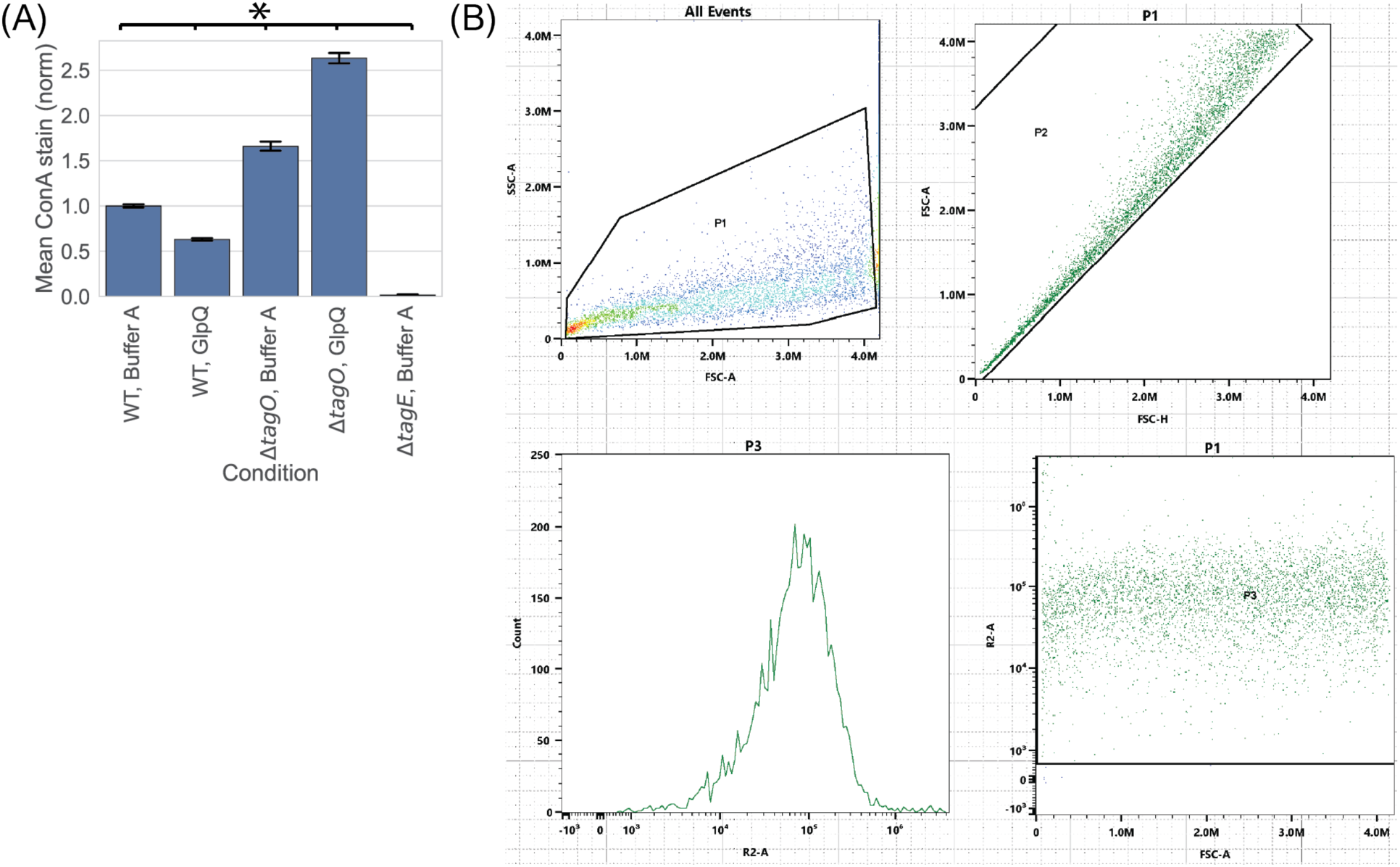
(A) Mean fluorescence intensity of Concanavalin A-AlexaFluor647 cell staining, following incubation either with PBS + Buffer A, or with PBS + GlpQ, measured by flow cytometry. GlpQ-treated cells show decreased Concanavalin A staining relative to PBS-treated cells, consistent with GlpQ enzymatically cleaving teichoic acids from the cell wall. Δ*tagO* cells show no such decrease in Concanavalin A staining upon GlpQ incubation, and Δ*tagE* cells lacking the enzyme responsible for teichoic acid glycosylation show negligible labeling. We observed increased Concanavalin A-staining in Δ*tagO* cells relative to wild-type that was amplified during GlpQ treatment, possibly due to the clumping phenotype of this mutant. Error bars show 95% confidence intervals across 3 biological replicates, 10,000 samples per replicate. All conditions show statistically significant differences as measured by one-way ANOVA followed by Tukey’s HSD post-hoc test, P<0.01. **(B)** Representative example of flow cytometry gating process used to generate data in (A).

**Figure S9:**
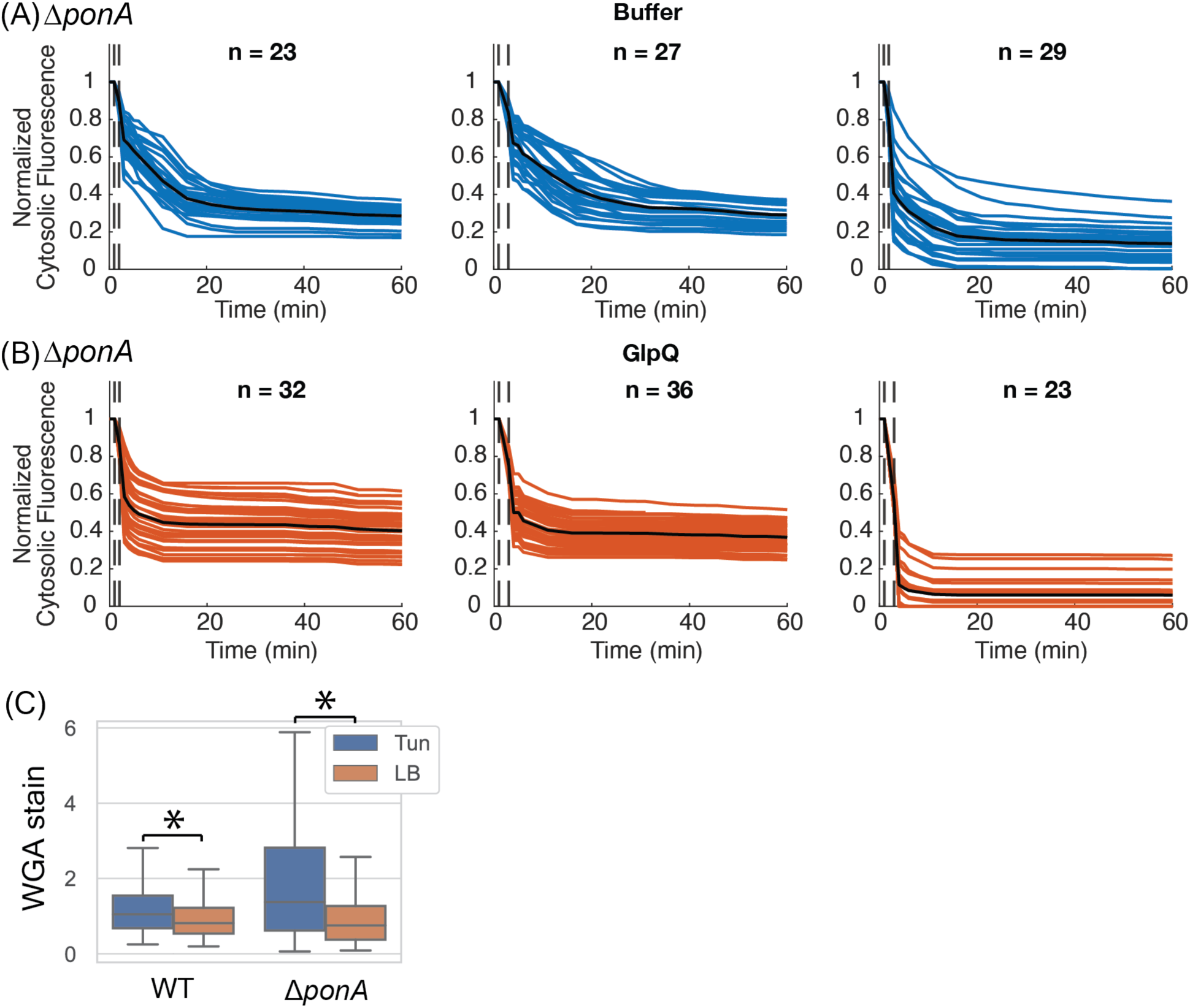
Ubiquitin-FlAsH immediately leaves the cytosol upon wall teichoic acid digestion and simultaneous lysis in Δ*ponA* cells. **(A-B)** Individual Ubiquitin-FlAsH fluorescent traces and experimental replicates following 2 min incubation with PBS +5% N-lauroylsarcosine and either **(A)** Buffer A alone, or **(B)** 40uM GlpQ. Cell counts are shown in figure captions. **(C)** Fluorescence of cells labeled with a wheat-germ agglutinin-AlexaFluor488 conjugate after incubation with LB alone or LB supplemented with tunicamycin for either 10 min (wild-type) or 20 min (Δ*ponA*). Wild-type LB: 538 cells. Wild-type tunicamycin: 271 cells. Δ*ponA* LB: 285 cells. Δ*ponA* tunicamycin: 288 cells. 3 biological replicates per condition. Significance tested by Student’s T-test, P<0.01.

**Figure S10:**
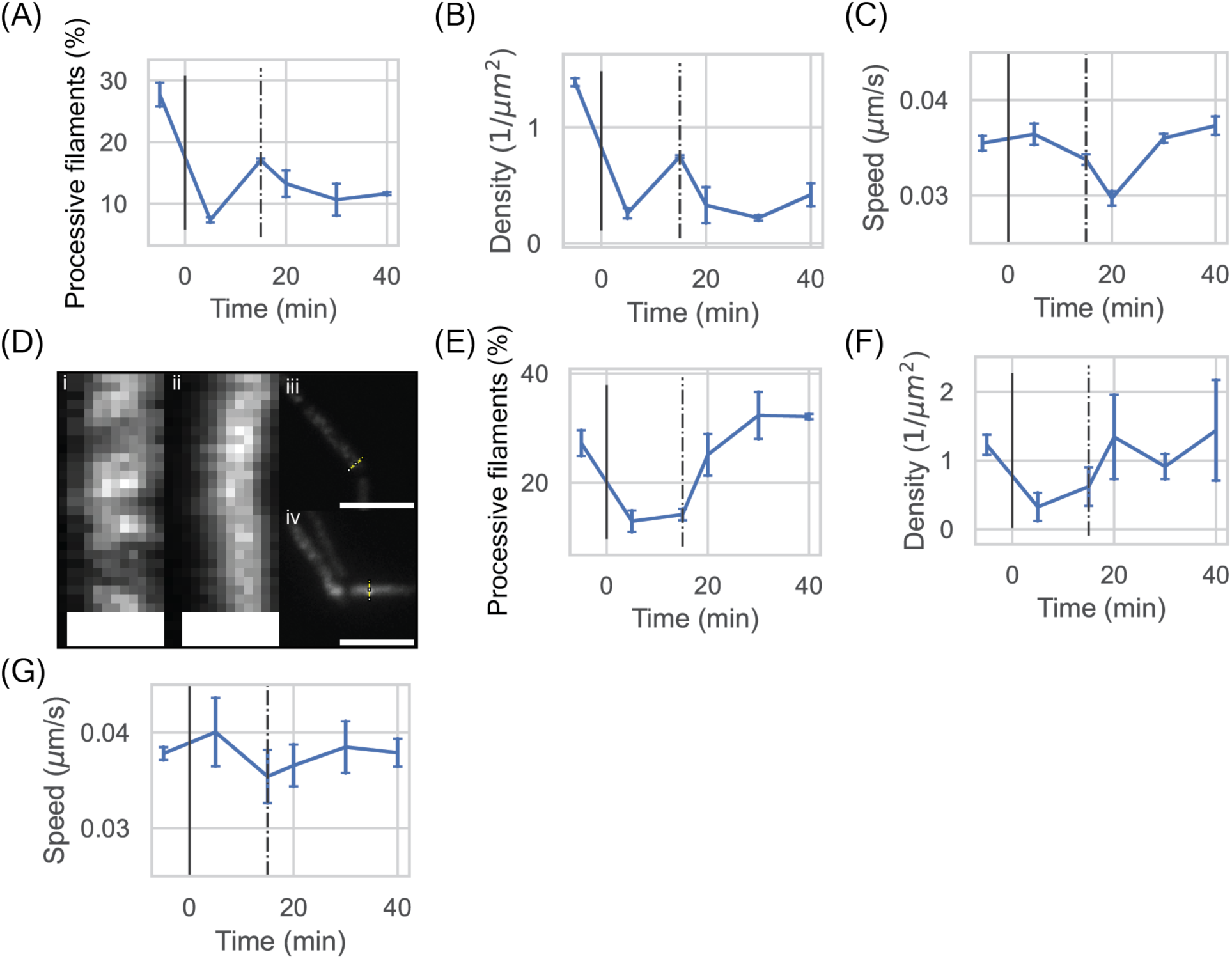
(A-C) Mbl-sfGFP activity time course during 25µM GlpQ treatment. Analysis performed on 10,976 discrete filament tracks from 2 biological replicates. **(A)** Percentage of processive filaments. **(B)** Spatial density of processive Mbl-sfGFP filaments. **(C)** Speed of processive Mbl-sfGFP filaments. All error bars show 95% confidence intervals across biological replicates based on bootstrap analysis. **(D)** Representative kymographs from TIRF imaging of (i) growing and (ii) non-growing wild-type cells 35 min after release from GlpQ treatment. Scale bars 1µm. (iii-iv) TIRF micrographs showing whole cells during a single timepoint of kymographs in (i-ii) respectively. Yellow lines show kymograph traces. Scale bars 5µm. **(E-G)** Mbl-sfGFP and MreB-mNeonGreen activity time course during PBS treatment. Analysis performed on 11,150 discrete filament tracks from 3 independent experiments. Data pooled from experiments with 15min and 20min PBS exposure. **(E)** Percentage of processive filaments. **(F)** Spatial density of processive Mbl-sfGFP filaments. **(G)** Speed of processive filaments.

**Figure S11:**
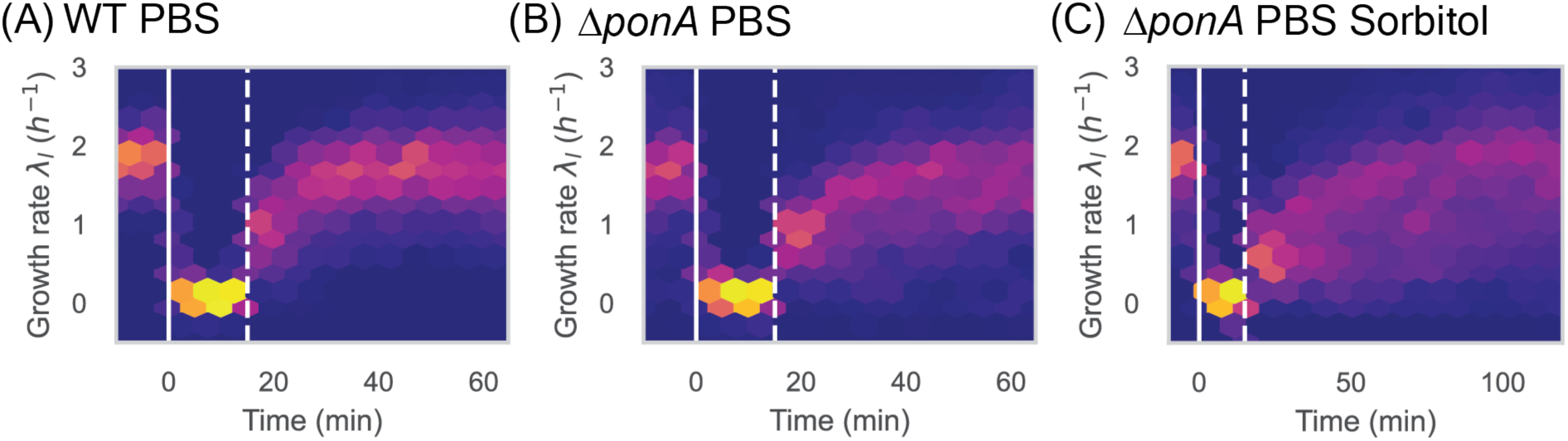
(A-B) Heat maps showing growth rate 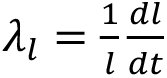 during recovery after incubation with either PBS + denatured GlpQ or GlpQ buffer for 15 min. **(A)** wild-type cells (3,830 discrete tracks, two independent experiments with denatured GlpQ, one with equivalent volume GlpQ buffer, no differences between conditions observed). **(B)** Δ*ponA* cells (2,142 discrete cell tracks, 3 biological replicates). **(C)** Heatmap showing Δ*ponA* cell length growth rate 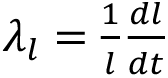 following 15-minute incubation with PBS + GlpQ Buffer (*not* GlpQ enzyme) (7,097 discrete cell tracks, 3 biological replicates), where exit from incubation is coupled to a 500mM Sorbitol hyperosmotic shock. Solid line shows onset of PBS incubation, dotted line shows exit into LB + Sorbitol.

**Figure S12:**
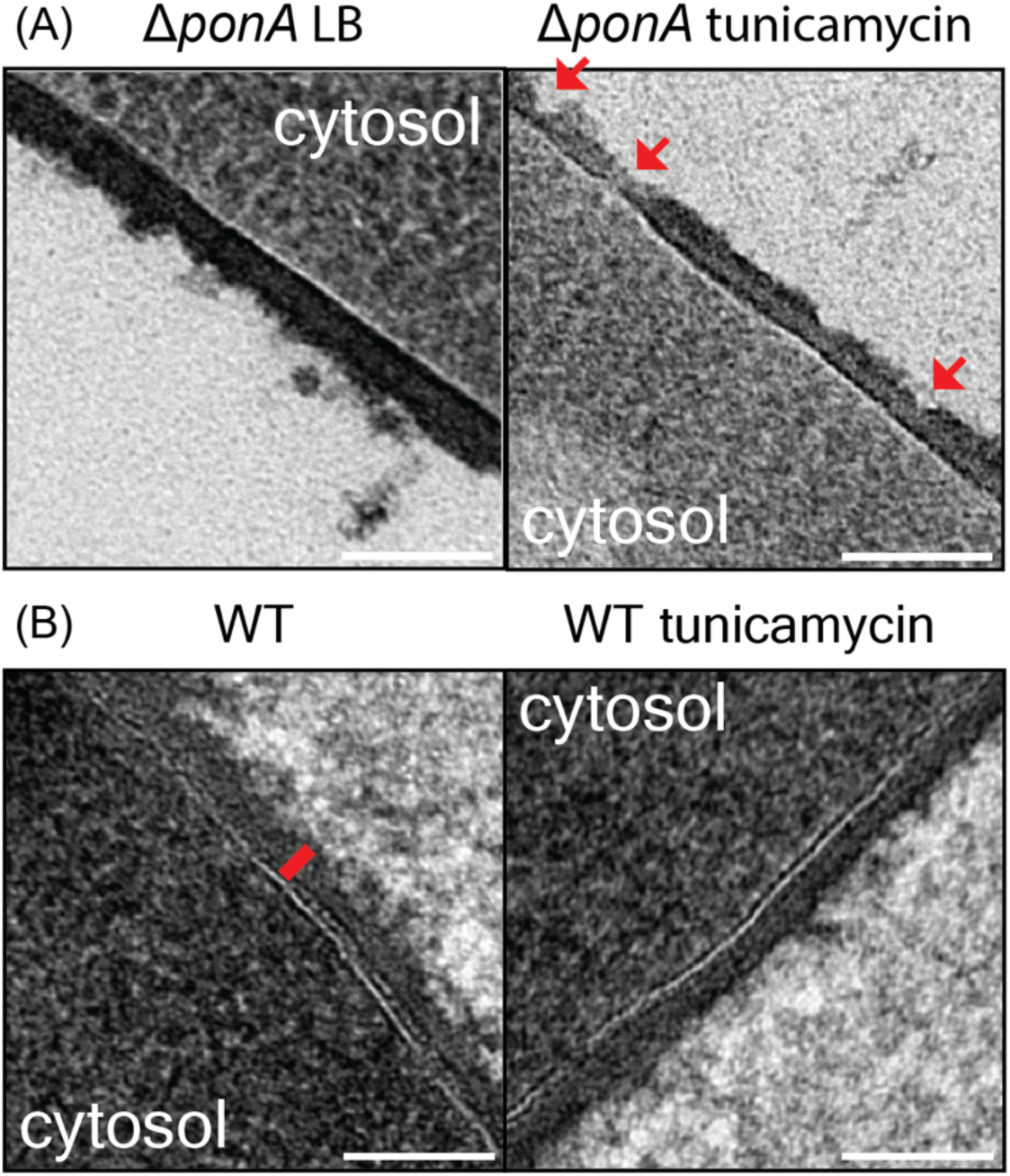
Transmission electron micrographs of the sidewall of exponentially growing cells grown in either LB or LB supplemented with 0.5𝜇g/mL tunicamycin for 30 min. Scale bars 100nm. **(A)** Δ*ponA* cells. Red arrows show sites of severe cell wall thinning during tunicamycin treatment. **(B)** Wild-type cells. Red bar shows the region used to calculate cell wall thickness, measured from the edge of the cell membrane (white line).

**Figure S13.**
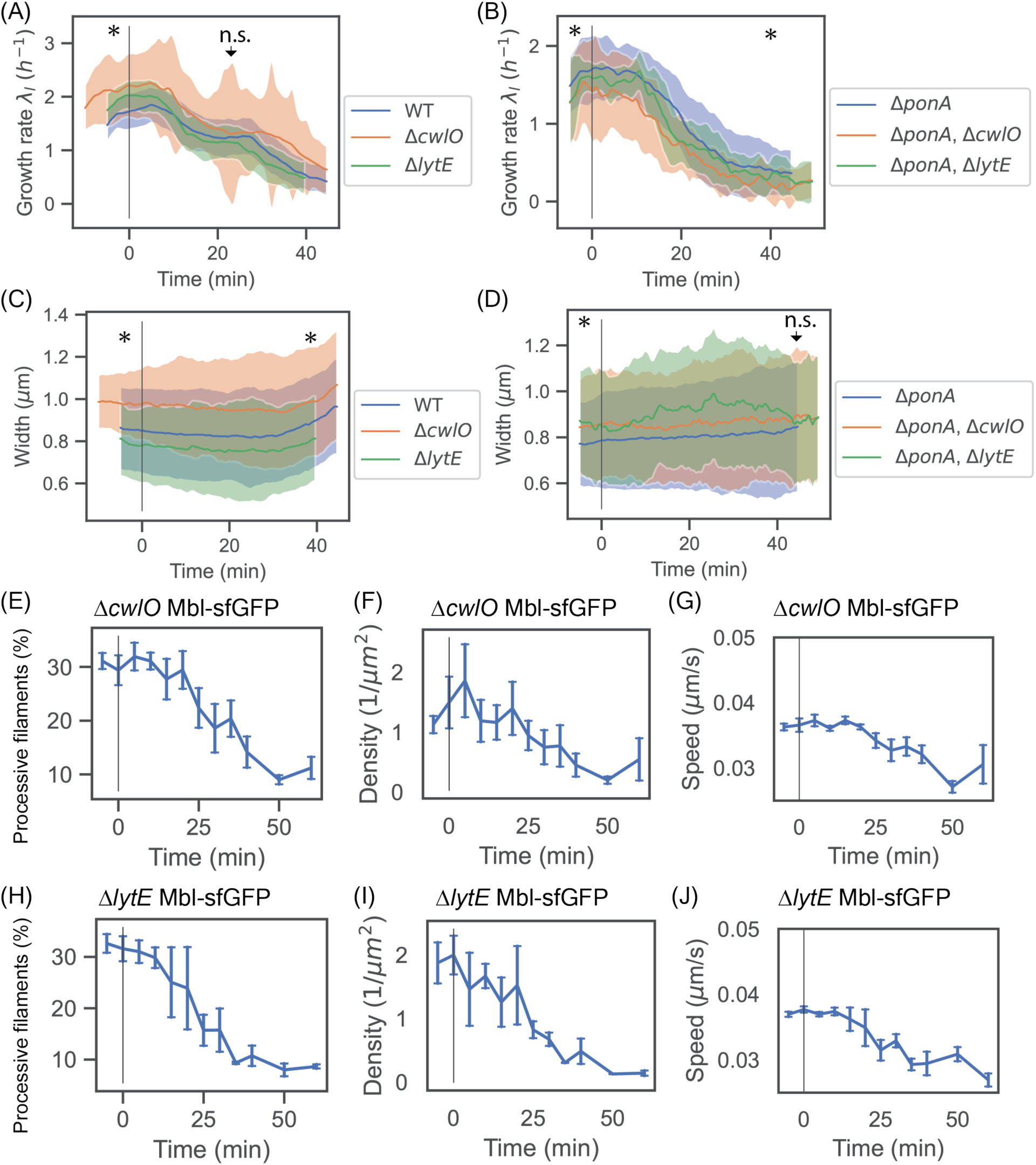
(A-B) Cell elongation rate during tunicamycin treatment for **(A)** wild-type, Δ*cwlO* and Δ*lytE* cells and **(B)** Δ*ponA*, Δ*ponA* Δ*cwlO* and Δ*ponA* Δ*lytE* cells. **(A)** Strains showed statistically significant differences in initial growth rate (time=-5min), but converged to a plateau growth rate that did not reject the null (equal) hypothesis (t=22 min following tunicamycin treatment). Significance calculated using Student’s T-test, P<0.01. Analysis performed over 1,091 discrete cell tracks from 5 experimental replicates (Δ*cwlO*), and 1,277 discrete cell tracks plotted from three experimental replicates (Δ*lytE*). Wild-type data replotted from Fig. 1B. **(B)** Hydrolase mutants showed significantly lower initial growth rates than the Δ*ponA* mutant (first measured timepoint), while the Δ*ponA* Δ*cwlO* mutant showed a significantly lower final growth rate than the Δ*ponA* mutant. All strains lacked the plateau in growth rate of wild-type cells. Analysis performed over 164 discrete cell tracks from two experimental replicates (Δ*ponA* Δ*cwlO*), and 105 discrete cell tracks plotted from two experimental replicates (Δ*ponA* Δ*lytE*). Δ*ponA* data replotted from Fig. 2A. **(C-D)** Cell width dynamics during tunicamycin treatment for **(C)** wild-type, Δ*cwlO* and Δ*lytE* cells and **(D)** Δ*ponA*, Δ*ponA* Δ*cwlO* and Δ*ponA* Δ*lytE* cells. **(C)** Strains showed statistically significant differences in width throughout, calculated using Student’s T-test, P<0.01 at the first and last measured timepoints. All strains display characteristic cell widening at long times. Analysis performed over 1,746 discrete cell tracks from 5 experimental replicates (Δ*cwlO*), and 2,432 discrete cell tracks from three experimental replicates (Δ*lytE*). Wild-type data replotted from Fig. 1B. **(D)** Hydrolase mutants showed statistically significant differences in width from Δ*ponA* mutant at the first measured timepoint, but showed no significant differences at the final timepoint (t=45 min tunicamycin exposure) and consistently lacked the characteristic cell widening of wild-type cells at long times. Significance calculated using Student’s T-test, P<0.01. Analysis performed over 1,747 discrete cell tracks from five biological replicates (Δ*ponA*), 632 discrete cell tracks from two experimental replicates (Δ*ponA* Δ*cwlO*), and 585 discrete cell tracks from two experimental replicates (Δ*ponA* Δ*lytE*). **(E-J)** Rod complex dynamics measured by tracking Mbl-sfGFP filaments during tunicamycin treatment for **(E-G)** Δ*cwlO* and **(H-J)** Δ*lytE* cells. **(E, H)** Percentage of processive filaments. **(F, I)** Spatial density of processive filaments. **(G, J)** Average speed of processive filaments. Analysis performed over 37,707 tracked filaments from four biological replicates (Δ*cwlO*) and 45,748 tracked filaments from three biological replicates (Δ*lytE*).

**Figure S14.**
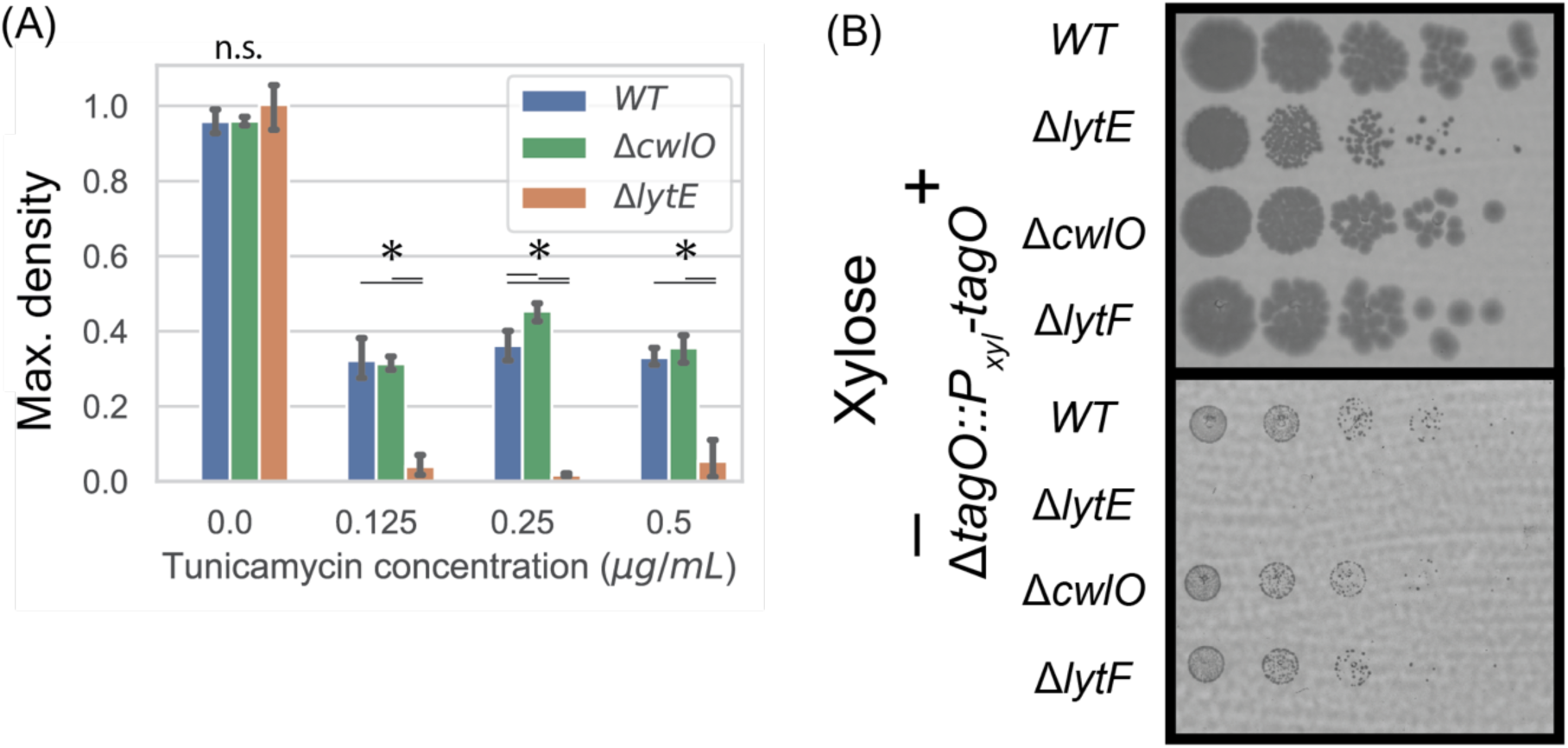
(A) Saturating culture density during growth in various concentrations of tunicamycin for wild-type, Δ*cwlO* and Δ*lytE* cells. Data shown from one representative biological replicate out of two, with four technical replicates per biological replicate. Error bars show 95% confidence intervals based on bootstrap analysis. Statistical significance tested with one-way ANOVA followed by Tukey’s HSD post-hoc test for different concentrations of tunicamycin, P<0.01. **(B)** Spot assays showing growth with and without *tagO* transcriptional inhibition in *P_xylA_-tagO* cells with the mutant backgrounds shown (two biological replicates).

**Figure S15:**
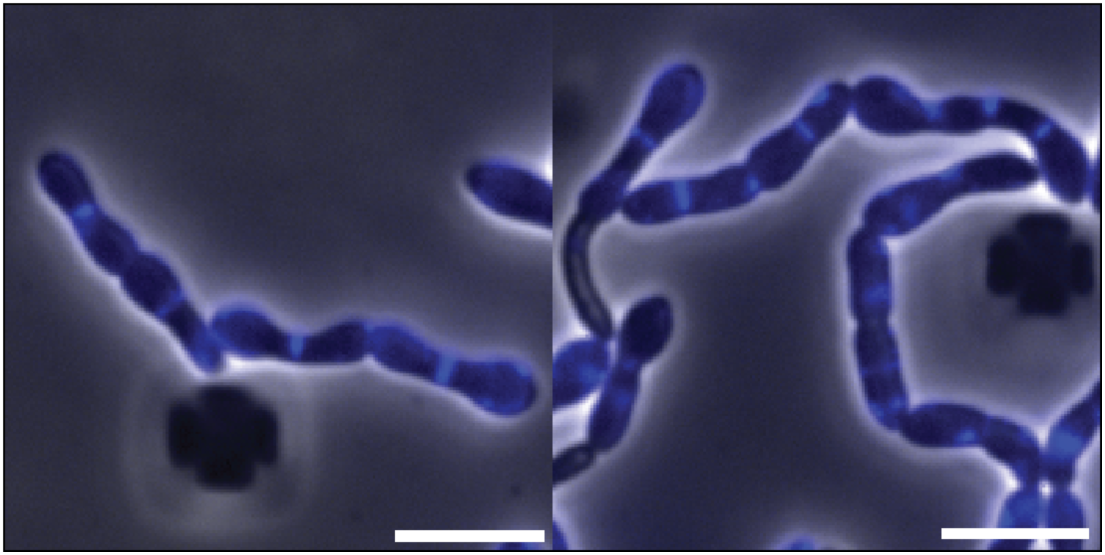
Cell bulging occurs gradually a “dumbbell” phenotype during teichoic acid depletion. Cell wall bulges become more prevalent with increasing distance from internal septa, presumably due to septal reinforcement of the local cell wall. Micrographs show HADA fluorescent staining in blue, superimposed over phase contrast images of cells following 45 min exposure to 0.5𝜇g/mL tunicamycin and 5min HADA incubation. Scale bars 5𝜇m. One biological replicate.

**Video S1:** 100X Phase timelapse of wild-type growth during 0.5 µg/mL tunicamycin treatment. tunicamycin exposure starts at 5min. Time step=20s. Scale bar 5µm.

**Video S2:** 100X TIRF imaging of Mbl-sfGFP motion in wild-type cells grown in LB. Time step=2s. Scale bar 5µm.

**Video S3:** 100X TIRF imaging of Mbl-sfGFP motion in wild-type cells grown in LB. Time step=2s. Scale bar 5µm.

**Video S4:** 100X TIRF imaging of Mbl-sfGFP motion in wild-type cells grown in LB, then treated with 0.5 µg/mL tunicamycin for 30min. Time step=2s. Scale bar 5µm.

**Video S5:** 100X TIRF imaging of Mbl-sfGFP motion in wild-type cells grown in LB, then treated with 0.5 µg/mL tunicamycin for 30min. Time step=2s. Scale bar 5µm.

**Video S6:** 100X TIRF imaging of Mbl-sfGFP motion in wild-type cells grown in LB, then treated with 0.5 µg/mL tunicamycin for 60min. Time step=2s. Scale bar 5µm.

**Video S7:** 100X TIRF imaging of Mbl-sfGFP motion in wild-type cells grown in LB, then treated with 0.5 µg/mL tunicamycin for 60min. Time step=2s. Scale bar 5µm.

**Video S8:** 100X TIRF imaging of PBP1-mNeonGreen dynamics during steady state growth, driven from a heterologous HyperSpank promoter with 10𝜇M IPTG in addition to the endogenous copy of *ponA*. Time step = 500ms. Scale bar 5µm.

**Video S9:** 100X TIRF imaging of PBP1-mNeonGreen dynamics, driven from a heterologous HyperSpank promoter with 10𝜇M IPTG in addition to the endogenous copy of *ponA*. Cells treated with 0.5 µg/mL tunicamycin for 90min. Time step = 500ms. Scale bar 5µm.

**Video S10:** 100X TIRF imaging of Mbl-sfGFP motion in Δ*ponA* cells grown in LB supplemented with 10mM MgCl_2_. Time step=2s. Scale bar 5µm.

**Video S11:** 100X TIRF imaging of Mbl-sfGFP motion in Δ*ponA* cells grown in LB supplemented with 10mM MgCl_2_, then treated with 0.5 µg/mL tunicamycin for 30min. Time step=2s. Scale bar 5µm.

**Video S12:** 100X TIRF imaging of Mbl-sfGFP motion in growing wild-type cells treated with 25uM GlpQ for 15 minutes. Timelapse taken 25 min post release into LB. Time step=2s. Scale bar 5µm.

**Video S13:** 100X TIRF imaging of Mbl-sfGFP motion in growing wild-type cells treated with 25uM GlpQ for 15 minutes. Timelapse taken 35 min post release into LB. Time step=2s. Scale bar 5µm.

**Video S14:** 100X TIRF imaging of Mbl-sfGFP in non-growing wild-type cells treated with 25uM GlpQ for 15 minutes. Timelapse taken 25 min post release into LB. Time step=2s. Scale bar 5µm.

**Video S15:** 100X TIRF imaging of Mbl-sfGFP in non-growing wild-type cells treated with 25uM GlpQ for 15 minutes. Timelapse taken 35 min post release into LB. Time step=2s. Scale bar 5µm.

