## Supplementary figures and images for "Wall teichoic acids regulate peptidoglycan synthesis by paving cell wall nanostructure"

### Video S1

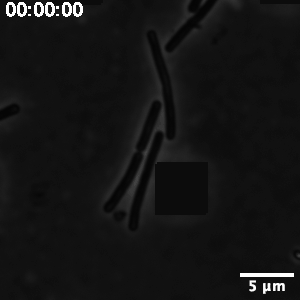

### Video S2

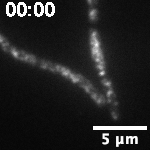

### Video S3

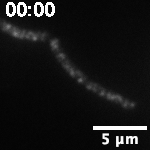

### Video S4

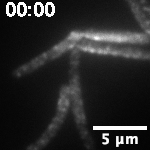

### Video S5

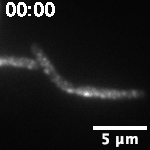

### Video S6

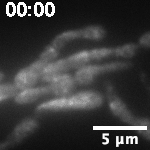

### Video S7

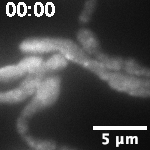

### Video S8

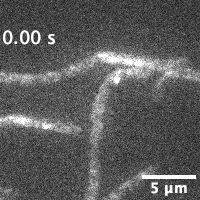

### Video S9

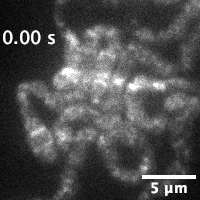

### Video S10

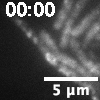

### Video S11

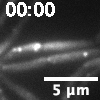

### Video S12

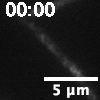

### Video S13

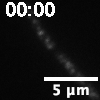

### Video S14

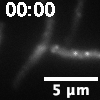

### Video S15

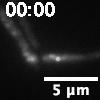
